# Spatial-temporal analysis of nanoparticles in live tumor spheroids impacted by cell origin and density

**DOI:** 10.1101/2021.10.26.465839

**Authors:** Aria Ahmed-Cox, Elvis Pandzic, Stuart T. Johnston, Celine Heu, John McGhee, Friederike M. Mansfeld, Edmund J. Crampin, Thomas P. Davis, Renee M. Whan, Maria Kavallaris

**Affiliations:** Children’s Cancer Institute, Lowy Cancer Research Center, UNSW Sydney, NSW, 2031, Australia; ARC Center of Excellence in Convergent Bio-Nano Science and Technology, Australian Center for NanoMedicine, UNSW Sydney, NSW 2031, Australia; School of Women and Children’s Health, Faculty of Medicine and Health, UNSW Sydney, NSW, 2031, Australia; Katharina Gaus Light Microscopy Facility, Mark Wainwright Analytical Center, UNSW Sydney, NSW, 2031, Australia; ARC Center of Excellence in Convergent Bio-Nano Science and Technology, Systems Biology Laboratory, School of Mathematics and Statistics, and Department of Biomedical Engineering, University of Melbourne, Parkville, Victoria, 3010, Australia; 3D Visualisation Aesthetics Lab, UNSW Art & Design, UNSW Sydney, NSW, 2021, Australia; ARC Center of Excellence in Convergent Bio-Nano Science and Technology, Monash Institute of Pharmaceutical Sciences, Melbourne, Victoria, 3052, Australia; School of Medicine, Faculty of Medicine Dentistry and Health Sciences, University of Melbourne, Parkville, Victoria, 3010, Australia; Precision Medicine, Australian Institute of Bioengineering & Nanotechnology, University of Queensland, QLD, 40679, Australia

**Author notes:** Corresponding author: Maria Kavallaris. Deceased (Edmund J. Crampin).

**Keywords:** Nanoparticles, tumor spheroids, microscopy, fluorescence imaging, mathematical modelling, uptake kinetics

## Abstract

Nanoparticles hold great preclinical promise in cancer therapy but continue to suffer attrition through clinical trials. Advanced, three dimensional (3D) cellular models such as tumor spheroids can recapitulate elements of the tumor environment and are considered the superior model to evaluate nanoparticle designs. However, there is an important need to better understand nanoparticle penetration kinetics and determine how different cell characteristics may influence this nanoparticle uptake. A key challenge with current approaches for measuring nanoparticle accumulation in spheroids is that they are often static, losing spatial and temporal information which may be necessary for effective nanoparticle evaluation in 3D cell models. To overcome this challenge, we developed an analysis platform, termed the Determination of Nanoparticle Uptake in Tumor Spheroids (DONUTS), which retains spatial and temporal information during quantification, enabling evaluation of nanoparticle uptake in 3D tumor spheroids. Outperforming linear profiling methods, DONUTS was able to measure silica nanoparticle uptake to 10 µm accuracy in both isotropic and irregularly shaped cancer cell spheroids. This was then extended to determine penetration kinetics, first by a forward-in-time, center-in-space model, and then by mathematical modelling, which enabled the direct evaluation of nanoparticle penetration kinetics in different spheroid models. Nanoparticle uptake was shown to inversely relate to particle size and varied depending on the cell type, cell stiffness and density of the spheroid model. The automated analysis method we have developed can be applied to live spheroids *in situ*, for the advanced evaluation of nanoparticles as delivery agents in cancer therapy.

---

Nanoparticles have been heralded for their potential to revolutionise cancer therapy, by improving drug delivery and reducing collateral toxicity of therapies in patients [1]. However, the diversity of their biophysical characteristics (e.g. size, shape, charge and surface coating) has also created challenges in attaining a robust understanding of how nanoparticles interact with the local and peripheral tumor environment, and has ultimately hindered their progression to the clinic [2–4]. Multidisciplinary studies to better evaluate how nanoparticle designs affect biocompatibility, circulation, extravasation and drug efficacy have been a key focus in recent years [5–8]. Yet, the quantification of nanoparticle tumor penetration has received less attention, and current analysis approaches are not optimized to account for cell and tissue variability. Attention in this area is arguably as critical as nanoparticle circulation and extravasation, as tumor penetration and subsequent cellular uptake will ultimately dictate nanoparticle efficacy.

A major variable in assessing nanoparticle uptake is the cell model that is used. The most common method to assess nanoparticle uptake is in two-dimensional cell culture where cells are grown on plastic dishes. However, these 2D systems do not recapitulate the cellular micro- or macro-environment of solid tumors, and thus cannot effectively model the barriers faced by nanoparticles to reach their intended cell populations in the human body. Consequently, research has shifted to three-dimensional (3D) cellular models such as tumor cells grown as spheroids [9,10]. This is because 3D spheroids have been shown to emulate key cellular parameters associated with solid tumors (tissue heterogeneity, cell mechanics, nutrient and oxygen gradients) and have been shown to model tumor growth and drug response in a more realistic manner than 2D cell cultures [11–13]. Tumor spheroids can also be augmented with additional cell types to add complexity and have unsurprisingly become a superior model to test fundamental nanoparticle characteristics [14–16]. However, current analysis of nanoparticle uptake in 3D spheroids has primarily relied on fixed samples, using a range of *ex situ* techniques such as, flow cytometry, sectioning and immunostaining, and transmission electron microscopy [10,17–19]. A major limitation with these methodologies is a lack of quantitative power which retains both kinetic and spatial information, hindering effective comparisons of nanoparticle kinetics in different spheroid models. Other methods have been employed with moderate success; for example, time-of-flight mass spectrometry, and *in situ* immunostaining, sectioning and subsequent microscopy [18,20–22]. Unfortunately, these methods are not always readily accessible, often require substantial downstream labor, time, and expertise, and continue to use static tissue samples (usually fixed or embedded), thereby losing valuable kinetic information.

Developments in confocal microscopy have enhanced the visualisation of nanoparticles in spheroids with improved spatial and temporal resolution, and greater depth and less phototoxicity [23–25]. It has been increasingly applied to study nanoparticle uptake in these complex cellular models, however, quantitative analysis has remained largely rudimentary, using limited timepoints and linear profiling methods [23,26,27]. Recently, paired correlation or models of diffusion have been applied to examine accumulation or dynamic nanoparticle correlation, however their application in 3D cellular models has been limited [24,28,29]. In depth imaging analysis of nanoparticle kinetics in 3D cellular models that mimic the tumor microenvironment is essential to understand the impact of biophysical characteristics of nanoparticle design on therapeutic delivery but may also reveal cell-dependent impacts on nanoparticle uptake which were previously unknown.

Here we examined the spatial and temporal quantification of nanoparticle uptake by developing an accessible and rigorous method to evaluate nanoparticle penetration and uptake kinetics in 3D spheroids. Using confocal time-course data of nanoparticle uptake in live tumor spheroids, we developed a quantitative and automated analysis pipeline for the <u>D</u>etermination <u>o</u>f <u>N</u>anoparticle <u>U</u>ptake in <u>T</u>umor <u>S</u>pheroids (DONUTS) which was used to determine how nanoparticle characteristics, in this case size, influence uptake kinetics into tumor spheroids of differing cellular origins *in situ*. This method is compatible with all major imaging platforms and is suitable for isotropic and anisotropic spheroids of varying cell types and complexity.

## RESULTS

### Spheroid models of glioblastoma and neuroblastoma show different patterns of nanoparticle uptake

To visualize nanoparticle uptake in live tumor spheroids, we first established spheroid models of glioblastoma (U87) and neuroblastoma (SK-N-BE(2)). Tumor spheroids were grown to sizes of approximately 500 µm in diameter at Day 3 (505 ± 18 µm and 494 ± 19 µm in U87 and SK-N-BE(2) respectively), as determined from spheroid growth curves (Supplementary Figure 1). This generated tumor spheroids which exhibited phenotypes of solid tumor growth including centralized cell death at the core and proliferative viability of peripheral cells (Supplementary Figure 2) [12,14]. Importantly, not only are the cancers from which these cell lines are derived of particular interest for nanoparticle delivery to improve patient outcome [30–32], but further, the tumor spheroids from these cell lines represent different cells of origin and show markedly different growth characteristics (Supplementary Figure 1). Brain cancer glioblastoma (U87) cell spheroids grew as regular (isotropic) neurospheres, akin to what has been reported in the literature [33,34], while in the pediatric peripheral nervous system cancer, neuroblastoma, (SK-N-BE(2)), spheroids grew with visual anisotropy and variability (Supplementary Figure 1).

To determine the impact of nanoparticle size on uptake, we selected a set of Sicastar®-redF silica nanoparticles (SiNP) of varying diameters (10, 30 and 100 nm) which met criteria for standardized reporting in bio-nano literature [35]. We initially validated that SiNP, which had no drug loading, showed no adverse effects on the viability of each cell line (Supplementary Figure 3, Supplementary Figure 4). As 2D cultures are often known to be more sensitive to drug exposures than 3D spheroids [11,12], we corroborated this null result using a 2D resazurin-based assay and showed no impact on cell viability up to 100 μg / mL and 72 hours exposure of U87 or SK-N-BE(2) cells with nanoparticles alone, or in combination with live-compatible membrane dyes (Supplementary Figure 3, Supplementary Figure 4). Together, these results provided a foundation for the investigation of nanoparticle uptake into these two different tumor spheroid models.

For the visualisation of nanoparticle uptake over time, we embedded glioblastoma (U87) or neuroblastoma (SK-N-BE(2)) spheroids in 1% low melt agarose, to reduce movement while retaining 3D structure and viability, and imaged SiNP uptake at 30 minute intervals for a total of 12 hours (Figure 1a). Confocal imaging demonstrated penetration of 30 nm silica nanoparticles into both glioblastoma and neuroblastoma spheroids (Figure 1). Interestingly, U87 glioblastoma spheroids appeared to exhibit lower uptake of SiNP (Figure 1b and Figure 1c) in contrast to SK-N-BE(2) neuroblastoma spheroids (Figure 1d and Figure 1e) over the 12 hour time-course. We also noted variable penetration of SiNP along the circumference of SK-N-BE(2) spheroids, likely related to the irregular shape of these 3D spheroids (Figure 1e), compared to U87 which showed consistent detectable fluorescence around the circumference of the spheroid (Figure 1c).

**Figure 1.**
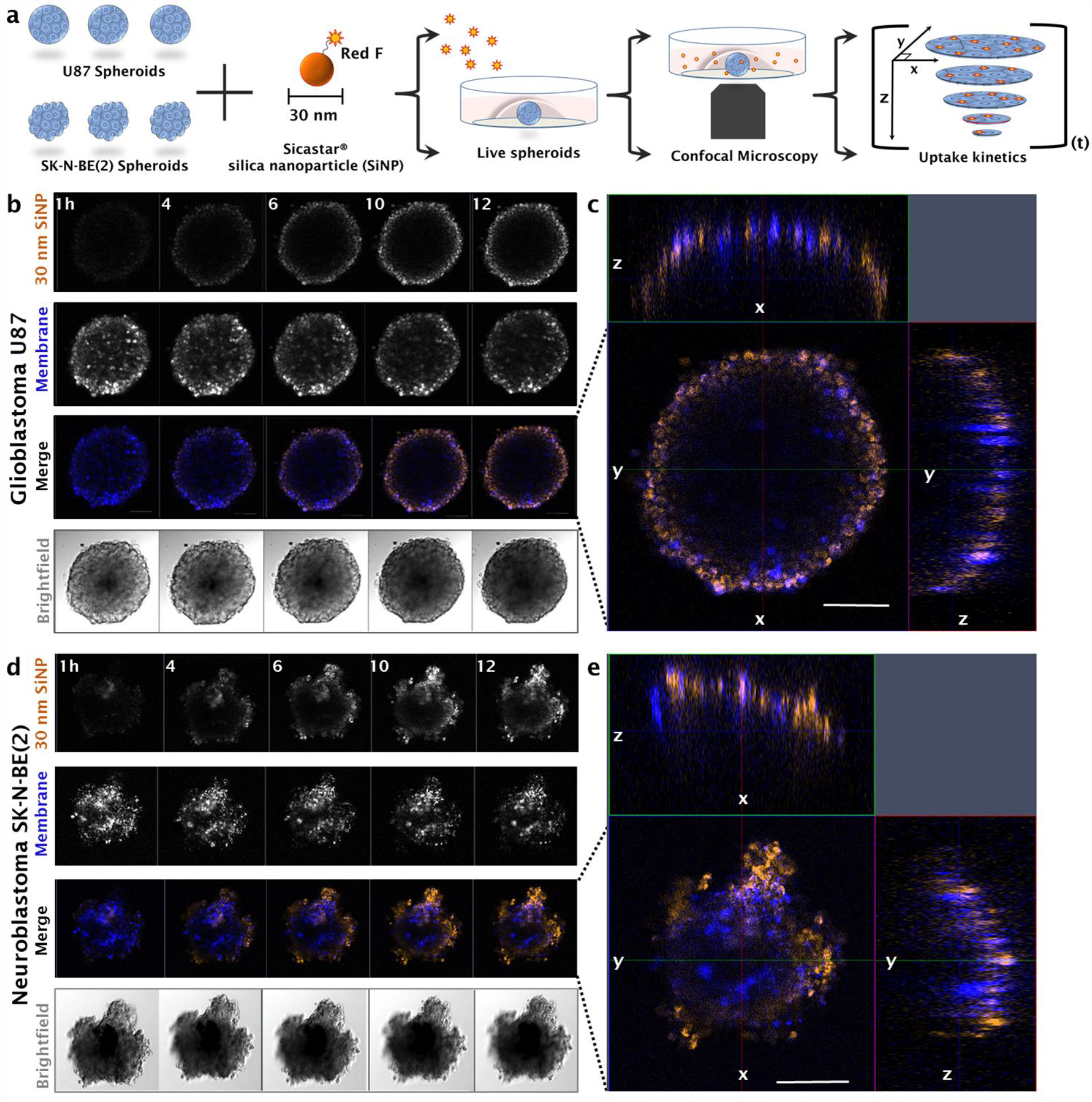
Visualisation of nanoparticle penetration into live glioblastoma (U87) and neuroblastoma (SK-N-BE(2)) tumour spheroids. **(a)** Graphical diagram outlining preparation of spheroids for live imaging of nanoparticle accumulation using confocal microscopy in XYZ over time (t). **(b)** Accumulation of 30 nm silica nanoparticles (SiNP, orange) uptake at 1, 4, 6, 10, 12 hours at the mid-plane (equator) of a glioblastoma (U87) spheroid labelled with a membrane dye (DiO, blue). **(c)** Representative image (n=3) of orthogonal data (XYZ) acquired at 12 hours post nanoparticle addition in U87. Scale bar, 100 µm. **(d)** Accumulation of 30 nm silica nanoparticles (SiNP, orange) uptake at 1, 4, 6, 10, 12 hours at the mid-plane (equator) of a neuroblastoma (SK-N-BE(2)) spheroid labelled with a membrane dye (DiO, blue). **(e)** Representative image (n=3) of orthogonal data (XYZ) acquired at 12 hours post nanoparticle addition in SK-N-BE(2). Scale bar, 100 µm.

### Development of a quantitative platform for the <u>D</u>etermination <u>o</u>f <u>N</u>anoparticle <u>U</u>ptake in <u>T</u>umor <u>S</u>pheroids (DONUTS)

After visualizing nanoparticle uptake in tumor spheroids, we turned to methodologies to analyse and evaluate this uptake in a quantitative manner. Initially, analysis was conducted using pre-established linear profiling methods to map fluorescence intensities along a defined vector in the mid-plane of the spheroid [23,26]. In this case we selected arbitrary angles for vectors at 0, 90 and 270 degrees (Figure 2a and Figure 2c in U87 and SK-N-BE(2) respectively). Fluorescence from SiNP could be detected at 150 µm from the core out to the circumference in U87 spheroids (Figure 2b) and showed similar intensity profiles across the three linear vectors selected. In contrast, SiNP appeared to penetrate deeper in SK-N-BE(2) spheroids (Figure 2c) which was confirmed by detection of fluorescence intensities above background at 50 - 100 µm from the core of SK-N-BE(2) spheroids (Figure 2d). In this case, the output from linear profiling was highly variable between different defined vectors, which introduced variability during analysis (Figure 2d).

**Figure 2.**
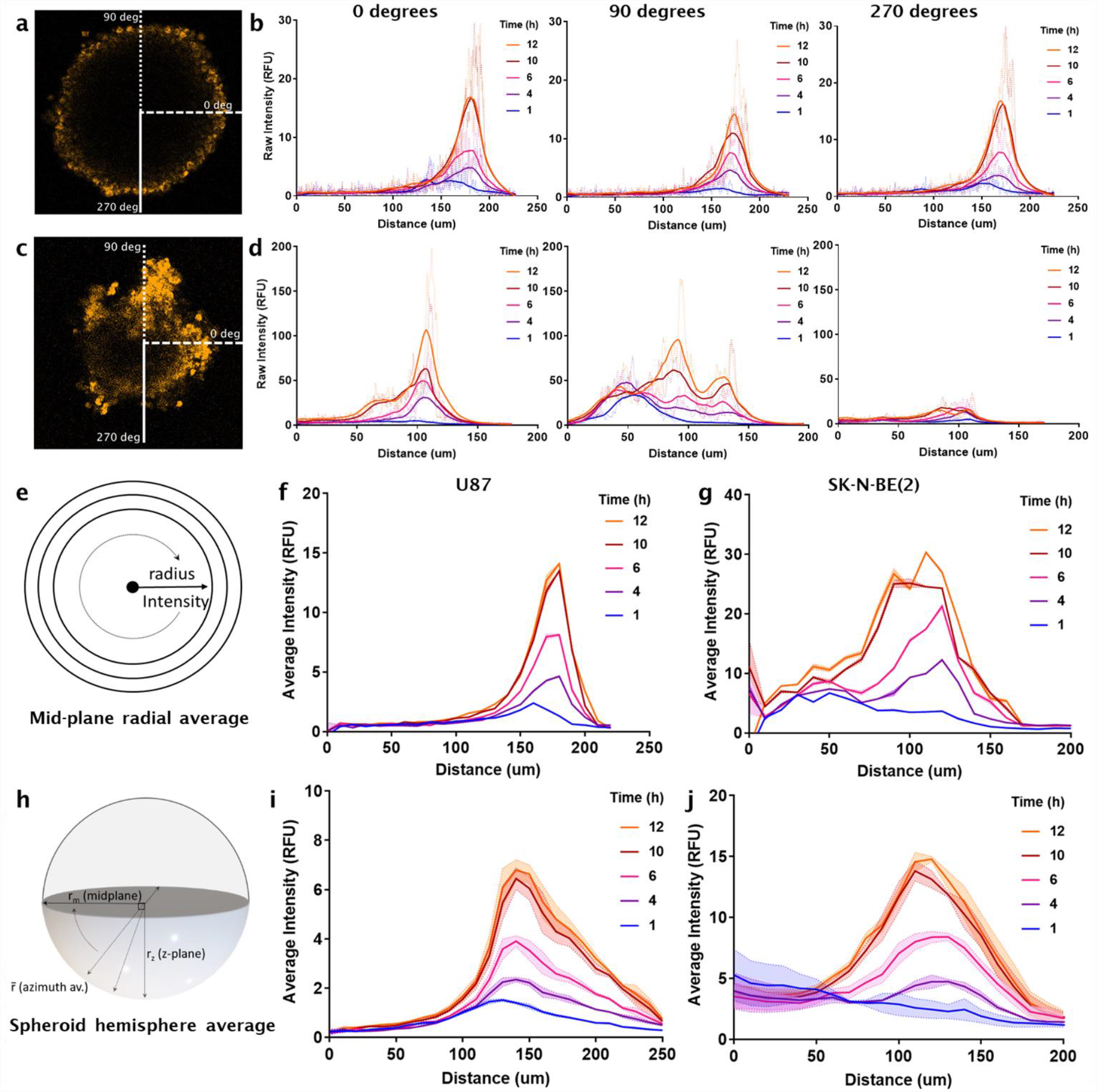
Quantification of nanoparticle uptake in tumour spheroids, from linear profiling to azimuth (hemisphere) averaging. **(a)** Representative images of 30 nm silica nanoparticle (SiNP) uptake in glioblastoma (U87) spheroid used for unidimensional linear profiling. **(b)** Plots of linear intensity of 30 nm SiNP uptake along three angles (0, 90 and 270 degrees) (dotted line, dashed line and solid line respectively in (a). **(c)** Representative mid-plane image of 30 nm SiNP uptake in neuroblastoma (SK-N-BE(2)) spheroid used for unidimensional linear profiling. **(d)** Plots of linear intensity of 30 nm SiNP uptake along three angles (0, 90 and 270 degrees) (dotted line, dashed line, and solid line respectively in (c). Dark solid lines, LOWESS regression fit imposed over raw data. **(e)** Graphical depiction of radial averaging of SiNP fluorescence across the mid-plane of the spheroids in (a) and (c). This was used to quantify nanoparticle uptake every 10 µm over time across mid-plane in **(f)** U87 and **(g)** SK-N-BE(2) over time (1 – 12 hours). **(h)** Graphical depiction of radial averaging extended in 3D (the azimuth average) to quantify nanoparticle uptake across spheroid hemispheres. **(i)** Representative azimuth quantification of SiNP uptake every 10 µm in U87 and **(j)** SK-N-BE(2) over time (1 – 12 hours). RFU, relative fluorescence units. Lines, mean of three analysis iterations. Shaded range, SEM.

To quantify uptake using all data available and thereby improve accuracy and reproducibility, nanoparticle fluorescence quantification was extended using a custom analysis platform in MATLAB (available in Supplementary Material) designed to calculate the average intensity along all defined radii in the mid-plane images (Figure 2a and Figure 2c), from the core of the spheroid to the spheroid circumference (Figure 2e). Using this script, we plotted average fluorescence intensities of SiNP at defined intervals (in this case, 10 µm). Automated iterative analysis on subsequent time-points then enabled the quantification of nanoparticle uptake over time in U87 and SK-N-BE(2) (Figure 2f and Figure 2g respectively).

To investigate whether this quantification changed when accounting for SiNP uptake throughout the spheroid, we extended our analysis to quantify nanoparticle penetration in 3D using all z-slice data. Analyses were therefore adjusted to calculate average nanoparticle fluorescence at a given radius, r, over all angles in 3D; termed the azimuth average (Figure 2h). This created the initial foundation for an automated and accessible analysis platform for the Determination of Nanoparticle Uptake in Tumor Spheroids (DONUTS) (full package and User Guide available in Supplementary Material). As part of DONUTS, users are prompted to load data, establish a binary mask to define boundaries of the tumor spheroid (using fluorescence data of labelled cells) and input basic parameters of experimental acquisition (pixel resolution, spheroid core, channel to be quantified etc.). In both cases in development (mid-plane and azimuth averaging), analysis required user defined selection of the core of the spheroid in XY or XY and Z. Selection was therefore repeated a minimum of three times per spheroid and the average of iterative results used for downstream analyses. Results from U87 (Figure 2i) and SK-N-BE(2) (Figure 2j) indicated good agreement between mid-plane averaging and 3D azimuthal averaging, with a larger average intensity of SiNP fluorescence in SK-N-BE(2) compared to U87 (maximum intensity values of 20 RFU versus 10 RFU respectively). More detail on the step-by-step prompts in DONUTS can be found in the Supplementary User Guide, which has been included to assist application of this methodology platform.

### Validation of azimuthal analysis using mathematically generated spheroid datasets

However, even isotropic tumor spheroids like U87 showed variations in azimuth quantification profiles between replicate uptake experiments (Supplementary Figure 5), which was more pronounced in SK-N-BE(2) which grew with visual anisotropy (Supplementary Figure 5).

To determine whether this variation was related to the independent and biological nature of spheroids, and to further validate DONUTS as a reliable methodology for measuring nanoparticle uptake, we generated simulated spheroids with differing densities of cell “objects” and modelled particle uptake into these convolved spheroids under different particle diffusion parameters (Figure 3). Parameters of particle diffusion used in simulations, included incremental probabilities of particle movement, which were also dependent on whether particles were intracellular (D(in)) or extracellular (D(out)) as they migrated through the simulated spheroids. Three separate simulations of the model were run with varying D(in) (0.0001, 0.001 and 0.01 µm^2^s^−1^). This model operated with the assumption that D(out) was orders of magnitude greater than D(in) which is a reasonable assumption given that particles should move more freely outside of cells than inside from spatial hindrance arguments alone. Results quantified using DONUTS demonstrated that particle diffusion through a “spheroid” with densely packed cell objects (High Density spheroid - HD) (Figure 3a) showed reduced penetration at low D(in) (0.0001 µm^2^s^−1^, Figure 3b) compared to particle diffusion through a Low-Density (LD) spheroid with small cells (SC) (Figure 3c) which, with increased space for extracellular diffusion (D(out)), showed greater penetration across different intracellular diffusion coefficients (Figure 3d). Further, adjustment of the intracellular diffusion coefficient (D(in)), a proxy for changing all active mechanisms of particle uptake and intracellular trafficking, had a stepwise impact on particle diffusion, where increasing D(in) increased the azimuth measurements of particle uptake into both models (Figure 3c and Figure 3d). A third spheroid (HDSC, Figure 3e) of high density like the HD spheroid (Figure 3a) but small cell (SC) objects like the LDSC spheroid (Figure 3c), was also investigated for penetration profiles that may appear intermediate due to equal density but an increased free surface area. Indeed, at low D(in) (0.0001 µm^2^s^−1^), profiles for the HDSC spheroid (Figure 3f) mimic that of the HD spheroid (Figure 3b), but as intracellular diffusion rates are increased, we observed greater penetration depth over time, approaching 50 µm from the core at D(in) = 0.001 µm^2^s^−1^, and detectable particles at the core at D(in) = 0.01 µm^2^s^−1^.

**Figure 3.**
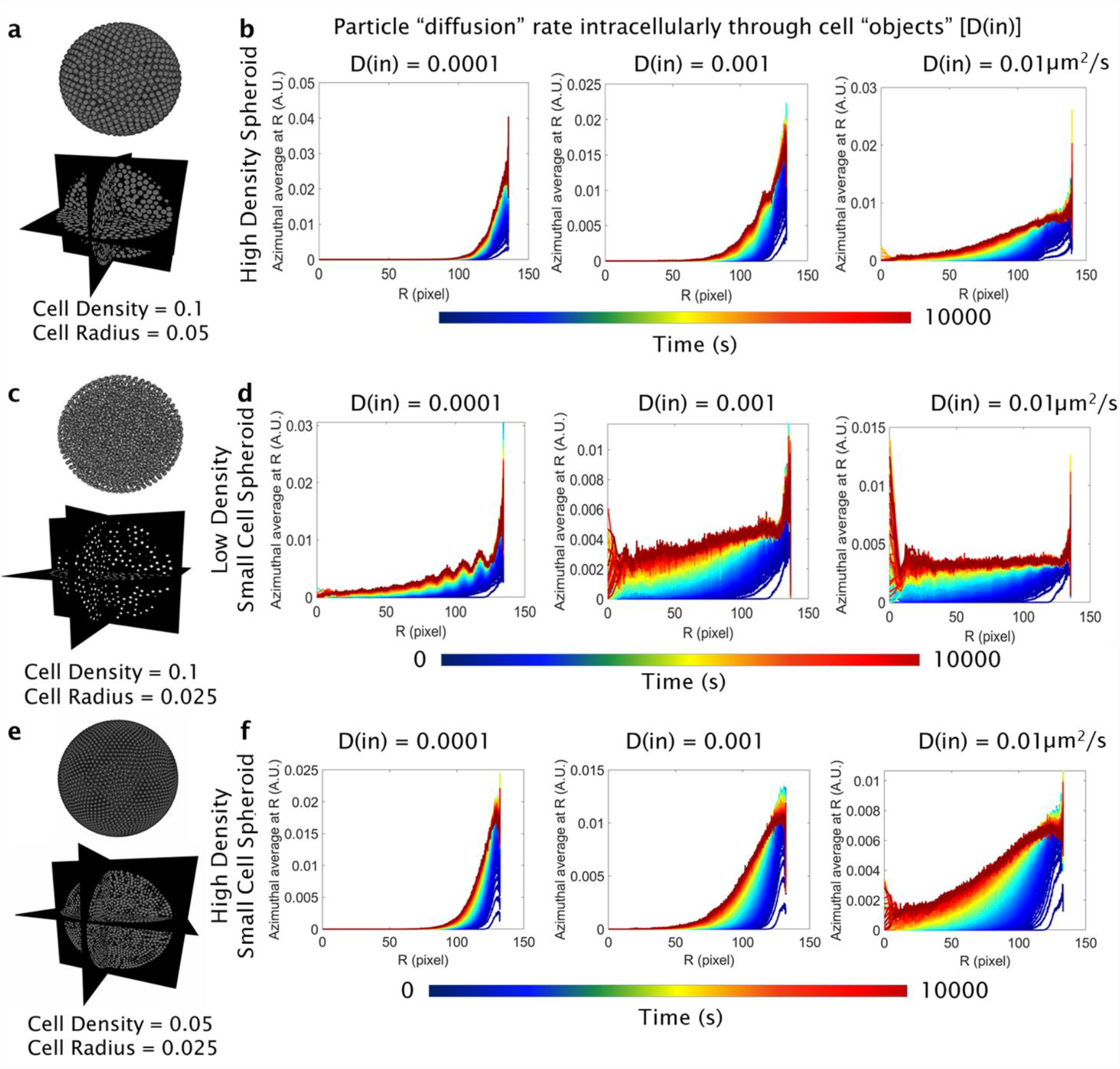
Simulated modelling of particle movement through convoluted spheroids of varying cell size and density using MATLAB (2020a, custom scripts). **(a)** “High density” (HD) convoluted spheroid of cell density 0.1 and cell radius 0.05 µm as a rendered 3D object, and orthogonal projection. **(b)** Simulated particle “diffusion” through the HD spheroid in (a) over time (10,000 s). Particle motion calculated at a fixed diffusion probability when moving extracellularly between cell objects, D(out) = 0.1 µm^2^s^−1^, and variable diffusion rates internally through cell objects, D(in) = 0.0001, 0.001, 0.01 µm^2^s^−1^. Analysis repeated for **(c)** “Low density Small Cell” (LDSC) convoluted spheroid of cell density 0.1 and cell radius 0.025 µm as a rendered 3D object, and orthogonal projection. **(d)** Simulated particle “diffusion” through the LD spheroid in (c) over time (10,000 s). Analysis repeated for **(e)** “High Density Small Cell” (HDSC) convoluted spheroid of cell density 0.05 and cell radius 0.025 µm as a rendered 3D object, and orthogonal projection. **(f)** Simulated particle “diffusion” through the HDSC spheroid in (e) over time (10,000 s). Pixel size = 0.1 µm.

### Quantitative *in situ* analysis of nanoparticle uptake in tumor spheroids are nanoparticle size and tumor model dependent

With our analysis methodology established, we next applied it to investigate the role of nanoparticle size (10 nm, 30 nm and 100 nm silica SiNP), in tumor spheroid accumulation [36]. To further strengthen our study, we incorporated a third cell model using non-small cell lung cancer (H460) tumour spheroids which were validated as above (See Supplementary data). Nanoparticle uptake was imaged in live spheroids at 30 minute intervals across 12 hours. Representative maximum intensity projections of SiNP uptake at 1, 4, 6, 10 and 12 hours post addition in U87 (Figure 4a), SK-N-BE(2) (Supplementary Figure 7) and H460 (Supplementary Figure 8) suggested greater accumulation and penetration of 10 nm SiNP compared to 30 nm and 100 nm SiNP (Figure 4, Supplementary Figure 7, Supplementary Figure 8). Orthogonal views confirmed retention of 3D shape in all spheroids (Figure 4b, Figure 4e, Figure 4g, Supplementary Figure 7, Supplementary Figure 8). Evaluation over the time-course (1 – 12 hours) suggested all three sizes of SiNP showed greater accumulation in SK-N-BE(2) spheroids, followed by U87 and then H460 spheroids. Of note, we observed variable fluorescence between independent biological replicates despite consistent imaging parameters (Supplementary Figure 6). This variability of fluorescence is likely a combined consequence of the fluorescence intensity and type of the individual nanoparticles, as well as biological interactions of nanoparticles in each cell line. Thus, quantification by this method alone has limitations.

**Figure 4.**
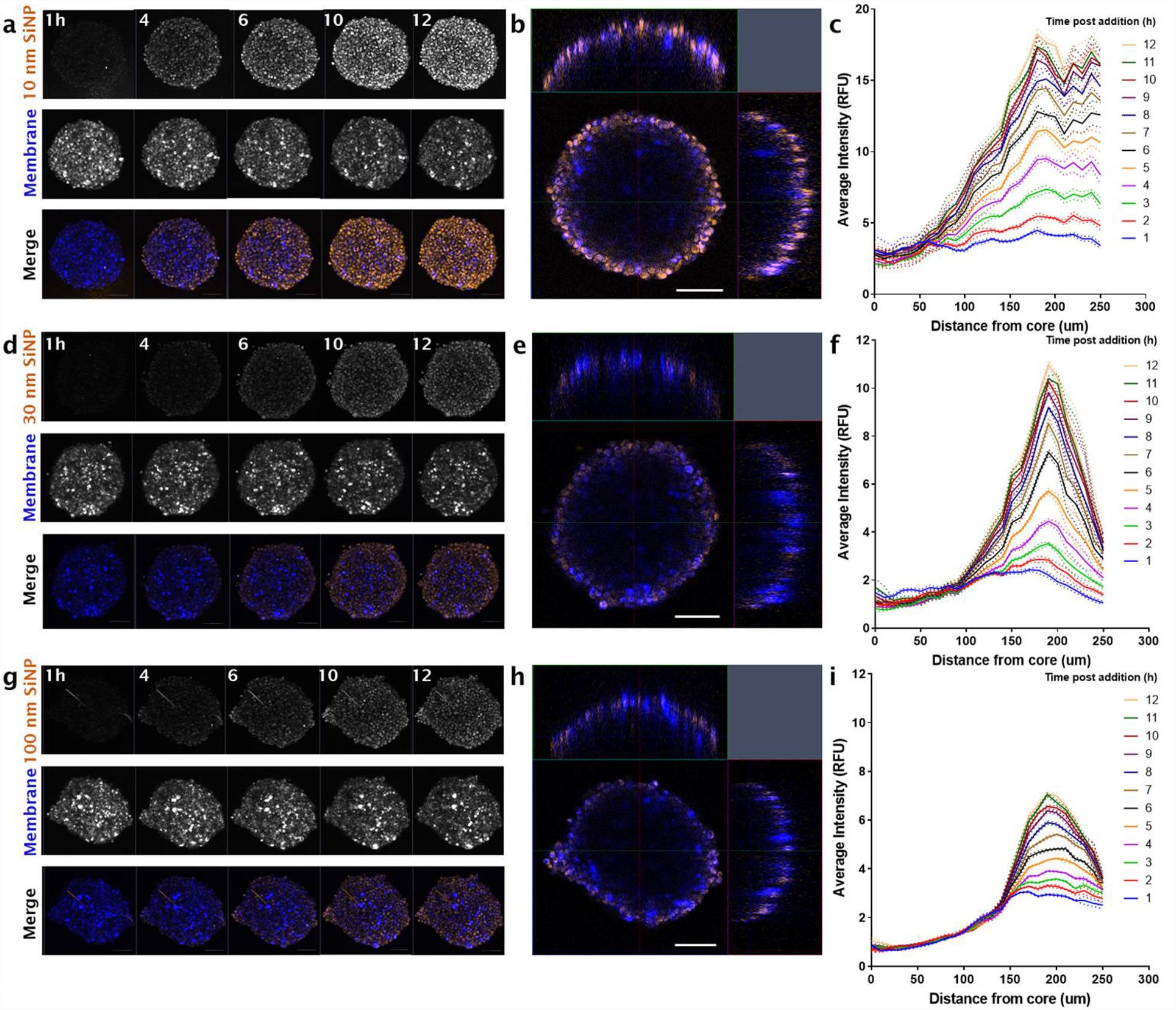
Silica nanoparticle (SiNP) uptake in glioblastoma (U87) tumour spheroids over time. **(a)** Representative maximum intensity projections of 10 nm SiNP, membrane (DiO) and merge at 1, 4, 6, 10, 12 hours post addition (Orange, SiNP; Blue, Membrane). Z-stack images acquired using a Zeiss 880 confocal microscope (Fast Airy, sequential frame-fast laser excitation at 488 nm and 561 nm, 10X objective). **(b)** Orthogonal (XY, XZ, ZY) merge of U87 spheroid at six hours post SiNP addition, representative of n = 3. **(c)** Representative quantification of nanoparticle uptake from the core of the spheroid to the circumference over time (1 – 12 h) with increased 10 nm SiNP penetration. Analysis conducted using a 3D azimuth averaging custom script, MATLAB (2020a). Workflow above was performed for 30 nm SiNP showing **(d)** maximum intensity projections over 1, 4, 6, 10, 12 hours post SiNP addition; **(e)** orthogonal merge at six hours and **(f)** azimuth quantification, respectively. Imaging and analysis also performed for 100 nm SiNP in panels **(g)** maximum intensity projections; **(h)** orthogonal merge at six hours and **(i)** quantification of 100 nm SiNP uptake. Lines, mean of n=3 analysis iterations. Dotted lines, SEM. Scale bar, 100 µm.

### Differences in nanoparticle penetration kinetics revealed via mathematical modelling

To evaluate whether silica nanoparticles did indeed show increased uptake in neuroblastoma (SK-N-BE(2)) spheroids compared to glioblastoma (U87) or NSCLC (H460) spheroids, we used the output of DONUTS azimuthal quantification to calculate the penetration kinetics (diffusivities), for each type of nanoparticle and tumor cell model considered. Penetration kinetics are independent to maximum fluorescence and used to transform the azimuth quantification into a rate of fluorescence change relative to distance through the spheroid. One approach to determine penetration kinetics is based on a forward-in-time, center-in-space (FTCS) diffusion model, often used to measure the mobilities of molecules in fluorescence recovery after photobleaching (FRAP) microscopy (Figure 5a) [37,38]. FTCS when applied in FRAP is a measure of the recovery of fluorescence inside a bleached area of a cell, which depends on the molecular motility of the fluorescent compound of interest and is one of the standard approaches to measure diffusion coefficients in live cell microscopy. In these experiments, we assumed that increases in fluorescence, due to nanoparticle penetration, within a sphere of radius *r* from spheroid center, would follow a similar trend to that observed in FRAP experiments. By monitoring the rise in average intensity within a sphere of radius *r* over time, we can extract the diffusivity at this radius from the spheroid center. Results demonstrated an average 10-fold increase in diffusion rates in 10 nm nanoparticles compared to 30 nm and 100 nm in our independent tumor spheroid models (Figure 5b and Figure 5d for U87 and H460 respectively). However, the diffusion coefficient from FTCS relies on multiple assumptions which inherently violated the biological dynamics we measured, including assumptions of a constant rate of particle diffusion (*i.e.*, that nanoparticle uptake kinetics do not change with radial distance) and assumptions of Gaussian fluorescence profiles, making it a simplified model of particle diffusion [38–40]. Evidence suggested that biological data deviated from this diffusion equation as nanoparticle fluorescence intensities dropped towards the core of both spheroid models, resulting in poor fit in diffusion kinetics <100-150 µm from the core (Supplementary Figure 9). This aligned with the large error ranges seen to the left of the dotted lines in Figures 5b – 5d, which corresponded with a drop in the adjusted R^2^ value between the fit of the data to the calculated FTCS diffusion coefficient (Supplementary Figure 9). This diffusion coefficient was also unable to distinguish differing diffusion rates of nanoparticles in irregular SK-N-BE(2) spheroids, despite clear trends in visual data and azimuthal quantification.

**Figure 5.**
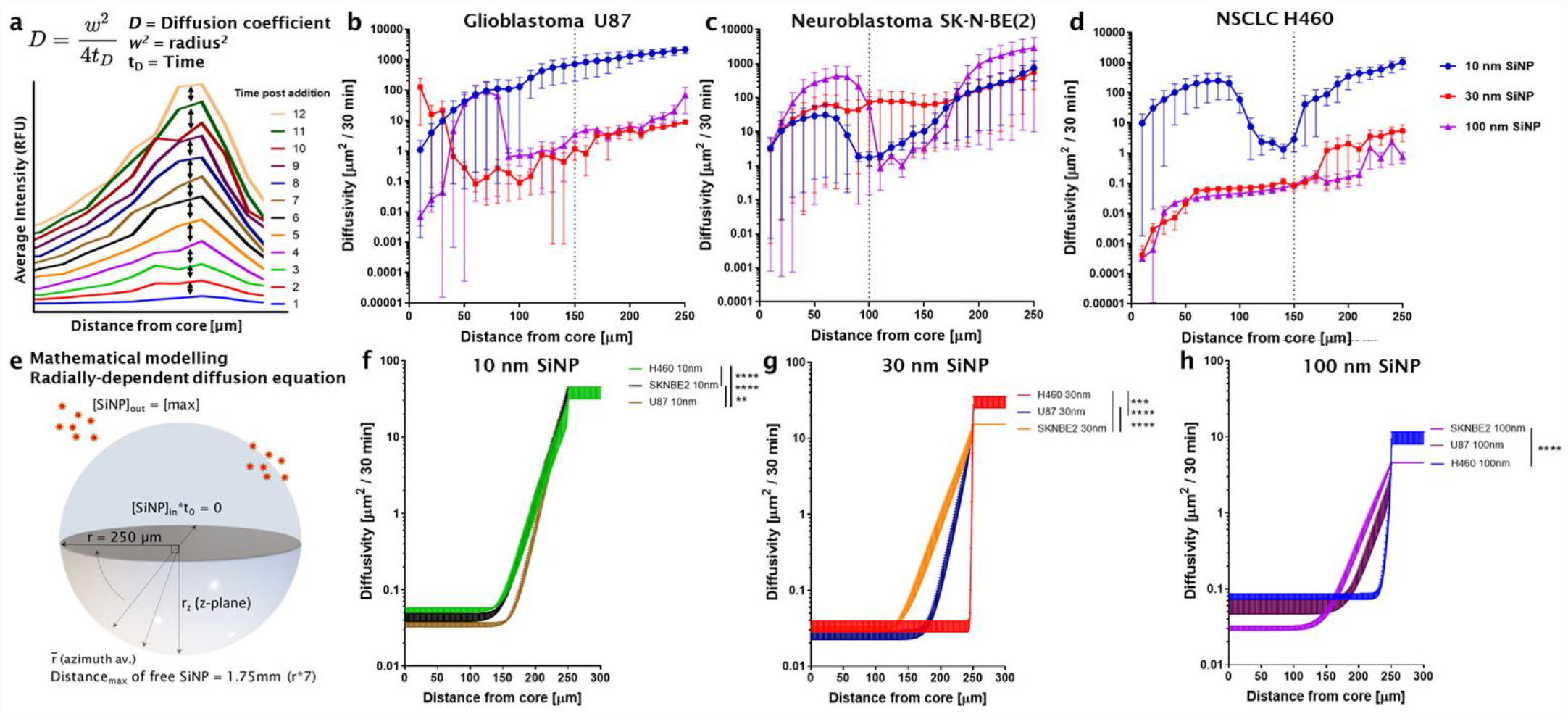
Calculation of penetration kinetics of silica nanoparticles (SiNP) in tumour spheroids. **(a)** Using output from 3D azimuth averaging, penetration kinetics of nanoparticles was quantified. Initial calculations used the diffusion coefficient based on a forward-in-time, center-in-space (FTCS) diffusion model. **(b)** Penetration kinetics of SiNP in glioblastoma (U87), **(c)** neuroblastoma (SK-N-BE(2)) and **(d)** non-small cell lung cancer (H460) calculated according to assumptions for a FTCS diffusion coefficient n=3 mean ± SEM. **(e)** Extension to a custom algorithm to model nanoparticle penetration kinetics. This was achieved by calibrating the numerical solution to a radially dependent diffusion equation to the fluorescence data in both space and time. **(f)** Penetration kinetics of 10 nm SiNP, **(g)** 30 nm SiNP and **(h)** 100 nm SiNP nanoparticles across U87, SK-N-BE(2) and H460 spheroids (n=3) using mathematical modelling and derived diffusion fitting. Significance between groups as indicated, ** *p* < 0.01, *** *p* < 0.001, **** *p* < 0.0001, Unpaired t-test with Welch’s correction.

To address this deviation and generate an improved kinetic equation for the data, a data-driven, mathematical model of diffusion was designed using MATLAB (script and guide available in Supplementary Material) and applied to quantify penetration kinetics across different nanoparticles and between different tumor spheroid models (Figure 5e). This method utilized the biological azimuth data as input and then calculated the numerical solution to a radially dependent diffusion equation in both space and time. Results confirmed a stepwise decrease in diffusivity as nanoparticle size increased, *i.e.*, 10 nm SiNP showed the greatest penetration kinetics compared to 30 nm and then 100 nm SiNP (Figures 5f - 5h).

Further, all three nanoparticles showed greater diffusivity in SK-N-BE(2) spheroids with penetration kinetic curves starting less than 150 µm from the spheroid core. This contrasted with U87 spheroids which showed penetration kinetic curves arising only above 150 µm from the spheroid core, indicative of negligible uptake below this distance. In contrast, nanoparticles showed differing trends in H460 spheroids, with a significant decrease in uptake kinetics of 30 nm compared to U87 spheroids, and both 30 nm and 100 nm particles when compared to the uptake kinetics in SK-N-BE(2) spheroids (Figure 5g and Figure 5h).

### Differing nanoparticle diffusion kinetics may be impacted by cell stiffness and spheroid densities

From simulations of nanoparticle diffusion in convolved spheroids (Figure 3), we observed that decreasing spheroid density resulted in increased particle penetration. Given trends of increased nanoparticle uptake in SK-N-BE(2) spheroids compared to U87 or H460 spheroids, we investigated whether this was associated with spheroid cell density, using spheroids which were chemically fixed and cleared, stained with DAPI and imaged using Lightsheet microscopy. We examined nuclei density of U87 using manual and automated nuclei detection (Figure 6a and Figure 6b). Automated nuclei detection was conducted using 3D feature finding [41] in MATLAB (representative in Figure 6c) to calculate nearest neighbor distances (NND). Automated detection of nuclei was then quality checked against manual counts across three z-stacks per spheroid, with tolerated variance defined at less than 7.5%. Nuclei were partitioned by distance from the core and showed a trend of increasing density towards the core of U87 spheroids, as measured by the reduced distribution of nearest neighbor distances (NNDs) between nuclei (Figure 6d). Interestingly, when the median cell diameter for U87 was overlaid on cell density data (Figure 6g), it appeared that cells within the tumour spheroid were more condensed, where the median distance between cells was in fact less than the median diameter of U87 single cells in suspension (19.05 ± 0.66 µm, Supplementary Table 1). The same analysis was applied to SK-N-BE(2) spheroids with manual (Figure 6e) and automated cell counts (Figure 6f), 3D feature finding (Figure 6g) and NND relative to median cell diameter (Figure 6h). As anticipated, distances between cells in SK-N-BE(2) spheroids were greater than the median cell diameter (12.16 ± 0.54 µm). Further analysis in H460 showed a similar trend to that of U87, with greater cell density where the NND of each neighboring cell was less than the median diameter of H460 cells in suspension (15.64 ± 0.11 µm) (Figure 6i, Figure 6j, Figure 6k and Figure 6l). However, the cell density of H460 spheroids, relative to the average cell diameter, was less than that of U87, indicating that U87 spheroids may have been expected to display the lowest nanoparticle uptake kinetics, if relying on spheroid density alone.

**Figure 6.**
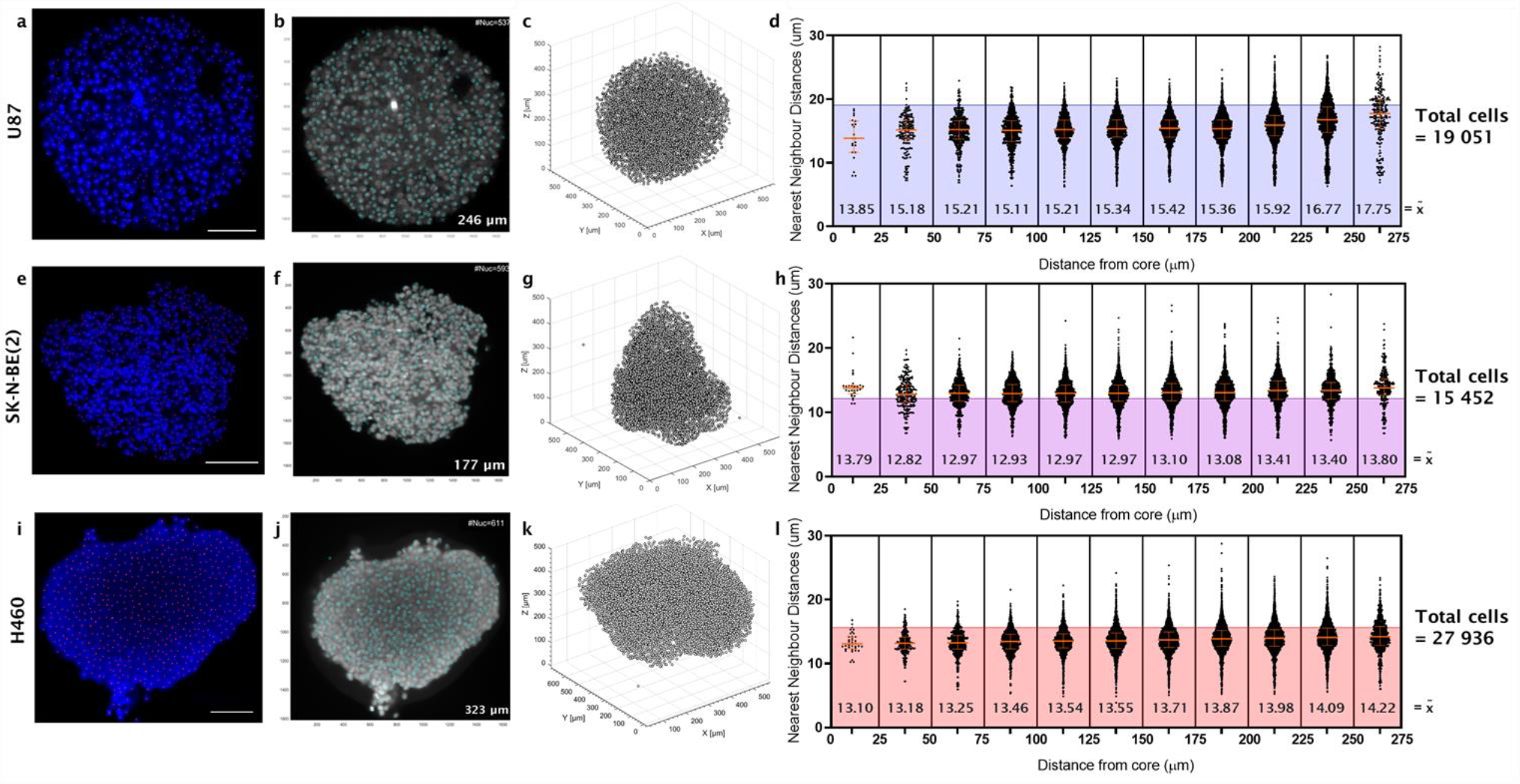
Density of nuclei in glioblastoma (U87) compared to neuroblastoma (SK-N-BE(2)) and NSCLC (H460) spheroids, prepared using optical clearing and imaged with lightsheet microscopy. **(a)** Glioblastoma (U87) spheroid which was fixed and optically cleared for nuclei localization using DAPI. Representative of n = 3. **(b)** Nuclei were detected using 3D feature finding in MATLAB (2020a, custom script adapted from [41]) and validated against manual counts in (a). **(c)** Nuclei coordinates in MATLAB were then used to calculate nearest neighbor distances (NND). **(d)** NND plotted at 25 µm intervals from the core to the circumference. This was repeated in neuroblastoma (SK-N-BE(2)) spheroids with **(e)** manual counts, **(f)** automated segmentation and **(g)** reconstructed spheroids for NND (n = 3) and **(h)** plotted NND. The same is then presented for NSCLC (H460) spheroids with **(i)** manual counts, **(j)** automated segmentation and **(k)** reconstructed spheroids for NND (n = 3) and **(l)** plotted NND. Scale bar in panels (a), (e) and (i), 100 µm. NND Plots in panels (d), (h) and (l): Orange lines, median with interquartile range across n=3 segmented spheroids per cell model. Shaded area behind data presents median cell diameter of cells in suspension (19.05, 12.16 and 15.64 µm, for U87 (blue) and SK-N-BE(2) (purple) and H460 (red) respectively). Median NND values (*x̃*) are given for every 25 µm interval from the spheroid core.

Finally, we investigated whether nanoparticle kinetics was associated with differences in the cell stiffness between the tumour spheroid models. Here, we employed force imaging cytometry to evaluate the stiffness and deformation of single cells from each tumour type (Supplementary Figure 10 and Supplementary Figure 11). Interestingly, H460 cells were significantly stiffer with a higher Young’s Modulus (1.56 kPa) compared to U87 (1.04 kPa) or SK-N-BE(2) (0.88 kPa) (*p* < 0.01, Supplementary Figure 10). H460 cells also had the lowest deformation potential (0.022 ± 0.001) versus SK-N-BE(2) (0.033 ± 0.004) or U87 cells, the latter which showed significantly higher deformation potential (0.056 ± 0.001) compared to H460 or SK-N-BE(2) cells (*p* < 0.01, Supplementary Figure 11). Collectively, this rigorous quantitative data analysis of nanoparticle uptake kinetics, using the DONUTS analysis platform, showed that nanoparticle penetration kinetics may be influenced not only by particle size, but also the cell density and stiffness of the tumor spheroid model.

## DISCUSSION

Here we developed an imaging and analysis approach for the Determination of Nanoparticle Uptake in Tumor Spheroids, termed DONUTS. This method does not require stable expression of a fluorescent marker protein in cells, immunostaining or IHC, and can be conducted using confocal microscopes with iterative or temporal data. It is designed to be accessible to users of broad disciplines (chemistry, biology, cancer research), capturing live cell uptake and compatible with various microscopy data formats and archival datasets. We applied DONUTS to investigate the impact of nanoparticle size on uptake and penetration kinetics of diverse 3D spheroid models, benchmarking an accessible and rigorous method to evaluate the effect of nanoparticle characteristics and cell model on penetration kinetics. Our findings revealed that nanoparticle uptake in live tumour spheroids was impacted by particle size, cell type, cell stiffness and density of the spheroid model.

As the current benchmark for *in situ* fluorescence measurements from visual data, linear profiling has benefits as an easily accessible, non-laborious analysis tool. However, quantification of nanoparticle uptake using this method was shown here to be heavily dependent on the vector selected, a known criticism of the technique [19,26,29]. Previous studies have moved to using multiple linear vectors to establish an intensity profile, however this adds manual labor and continues to exclude spatial data [29]. Variation in results by linear profiling methods may be attributed to the variability in fluorescence in any given z-plane, or an influence of the biological model itself, such as spheroids which deviate from isotropy as we have shown here in the neuroblastoma SK-N-BE(2) spheroids. In an isotropic spheroid model, such as glioblastoma (U87), our data suggests it may be sufficient to quantify nanoparticle uptake in a single z-plane, on the proviso that the plane selected is the equatorial (mid-)plane of the spheroid. U87 cells are well regarded as models which form tightly packed and isotropic spheroids [33,34] which was ideal for establishing our analysis platform. In contrast, in SK-N-BE(2), and in primary patient tumor samples, this same isotropy was not observed when cells were grown as spheroid cultures [42–44]. This can become increasingly common with the addition of multiple cell types (fibroblasts, endothelial cells, macrophages) and the formation of complex tumor organoids [44–46], necessitating analysis that incorporates all spatial and temporal information, that DONUTS provides.

Additionally, irrespective of isotropy, the outcome of linear profiling is the same: namely that much of the visual data acquired is discarded, and objective comparisons of different nanoparticles in different spheroid models become difficult. This undermines the applicability of linear profiling for effectively and reliably comparing nanoparticle designs for improved tumor penetration and uptake. This has been supported by previous studies which have highlighted the importance of a higher sample number to ensure reliability or moving to kinetic quantification as a means for more robust spatial analysis [21,24,29]. However, increasing sample number can quickly become labor- and cost-intensive for the in-depth quantification and comparison of nanoparticle penetration and uptake kinetics within these tumor models.

Thus, we focused on an *in situ* and automated method to evaluate nanoparticle uptake within spheroids, which retains temporal resolution and incorporates all accessible spatial information [19,47,47]. In establishing DONUTS, the application of automated, 3D radial averaging (azimuth averaging) enabled objective quantification of nanoparticle uptake in live spheroids over a 12-hour time course with single cell spatial resolution (10 µm, from the core to the circumference) and demonstrated the capacity of DONUTS to quantify nanoparticle penetration *in situ*, without any bias of manually defined vectors.

While DONUTS enabled effective quantification of nanoparticle uptake in our diverse tumor spheroid models, ultimately evaluation between models and different nanoparticles was still influenced by variations in fluorescence intensity profiles. Thus, we applied the output of DONUTS as the input data for calculating nanoparticle kinetics, effectively normalizing fluorescence for a direct comparison of nanoparticle uptake between spheroid models. Calculating the penetration kinetics of nanoparticles in cancer spheroids or organoids is an important step towards robust evaluation of nanoparticle uptake and falls under the drive toward consistent and reproducible reporting in bio-nano literature, for the direct comparison of nanoparticle designs between different studies [19,35]. We further validated the nanoparticle kinetic findings by incorporating mathematical modelling into our analysis, which has previously been used to investigate how nanoparticle characteristics influence cell uptake, or how tumor spheroid development may impact drug delivery, both in a high throughput and iterative manner [48–51]. Mathematical modelling independently validated DONUTS quantification of particle uptake with known diffusion properties in simulated spheroids of differing cell densities. We also applied a computational model to our analysis of nanoparticle uptake kinetics, using a model of radial-dependent nanoparticle diffusion informed by biological data. This generated enhanced fit and kinetic curves which were used to directly evaluate the uptake of different nanoparticles in tumor models of independent and diverse cell types. This has the capacity to be expanded to facilitate rapid analysis of biologically complex tumor models and drug loaded nanoparticles of varying designs. Our modelling confirmed that an increase in nanoparticle penetration inversely correlated with nanoparticle size, where 10 nm silica particles showed the greatest uptake kinetics compared to 30 nm and 100 nm particles, respectively. While the influence of nanoparticle size on uptake is well accepted [52–55], we also demonstrated that nanoparticle penetration in 3D appeared to be tumor model dependent. Silica nanoparticles showed greater uptake in SK-N-BE(2) compared to U87 and again to H460 spheroids at all particle sizes, suggesting the spheroid model has intrinsic properties which will influence nanoparticle delivery efficiency. Glioblastoma and NSCLC spheroids were highly compact, such that measurements of free space between adjacent nuclei were negligible, and showed evidence of cell crowding [43,56], potentially resulting in changes in cell size towards the core. In contrast, neuroblastoma spheroids held consistent and detectable free space throughout the model, which agreed with simulated low density spheroid data and may have contributed to increased nanoparticle uptake kinetics. We also showed that NSCLC cells had a greater stiffness and lower deformation potential compared to glioblastoma or neuroblastoma; characteristics of solid tumors which have been shown to hinder nanoparticle delivery in the past [3,57–60]. While we identified changes in cell stiffness and cell density in these spheroid models, there are of course several mechanisms of uptake (including active transport and transcytosis) which may also contribute to differential nanoparticle penetration kinetics and can now be evaluated using these methodologies in future studies [60]. Our analysis also identified differences in nanoparticle kinetics at the core of both spheroid models. A stark reduction in diffusivity of silica nanoparticles occurred at approximately 100 - 150 µm from the spheroid core for the 30 and 100 nm particles. Penetration depth of nanoparticles is an under studied measurement which is clinically relevant, as penetration and subsequent cell uptake will ultimately dictate nanoparticle efficacy [61]. For instance, Manzoor *et al*, demonstrated that Doxil has a penetration distance of less than 20 µm which could be increased to 78 µm with mild hyperthermia for enhanced efficacy [62]. Further, numerous studies have used fluorescence data of penetration to support the progression of nanoparticle designs for further clinical testing [10,26,52,63].

## CONCLUSION

Our study has shown that the spatial and temporal analysis of nanoparticle uptake kinetics are impacted by cell type, cell stiffness and density of the spheroid model. This was achieved by our development of a quantitative analysis tool to effectively evaluate the impact of nanoparticle characteristics on penetration kinetics into live tumor spheroids *in situ*. This method provides an accessible and robust quantitative platform, complete with proof-of-concept study and user support documents that will be valuable to facilitate analysis and advance the understanding and development of nanoparticle designs for enhanced clinical translation.

## METHODS

### Nanoparticles

Sicastar®-F red fluorescent silica nanoparticles were purchased from Micromod Partikeltechnologie GmbH (Germany) in sizes of 10 nm, 30 nm, and 100 nm with unmodified surface coatings. Specifications have been included in supplementary documents. Peak fluorescence excitation and emission for these particles are reported to be 569 / 585 nm, respectively. Fluorescence spectra and size distributions were confirmed using fluorometry on a Synergy Neo2 HTS Multi-Mode Microplate Reader (BioTek, USA) and DLS on a Zetasizer Nano (Malvern Panalytical, UK), respectively.

### Cell culture

U87 glioblastoma (ATCC HTB-14) and SK-N-BE(2) neuroblastoma (ATCC CRL-2271) were cultured in Dulbecco’s Modified Eagles Medium (DMEM) (Sigma-Aldrich, Australia) supplemented with 10% fetal bovine serum (FBS) (Life Technologies, Australia). H460 NSCLC (HTB-177) were cultured in Roswell Park Memorial Institute (RPMI) media (Sigma-Aldrich, Australia) supplemented with 10% fetal bovine serum (FBS) (Life Technologies, Australia). Cells were passaged at 70 – 90% confluency using 1% PBS and trypsin/EDTA (0.25%,0.02%) in T75 tissue culture flasks (Merck, Australia) and cultured at 37°C, 95% humidity and 5% CO_2_. Cells were cultured to a maximum of 32 passages or three months. For SK-N-BE(2) as a semi-adherent cell line, suspended and adherent cells were collected. All cell lines were tested regularly and found to be free of mycoplasma.

### Cell viability studies

Toxicity of Sicastar® nanoparticles was investigated in 2D cell culture using a modified Alamar Blue cytotoxicity assay [64]. In brief, glioblastoma U87 and neuroblastoma SK-N-BE(2) cells were seeded at 2 x 10^3^ cells / 100 uL / well in transparent 96-well plates (Interpath Services, Australia) in DMEM + 10% FBS. NSCLC H460 cells were seeded at 7 x 10^2^ cells / 100 uL / well in RPMI + 10% FBS. After 24 hours, silica nanoparticles were added at concentrations of 1, 10 or 100 ug / mL with or without membrane dye, DiO (1 uM). Doxorubicin (Sapphire Bioscience, Australia; 0.1 - 0.25 uM) was used as a positive control for the assay. After 72 hrs, 20 uL of resazurin blue reagent (Sigma) was added to each well and incubated for a further 6 – 12 hrs for reduction by active mitochondria, before spectrophotometry at 470-495nm using a Benchmark Plus Microplate Reader (BioRad, USA). 3D toxicity of silica nanoparticles was investigated in glioblastoma, neuroblastoma and NSCLC spheroids using a CellTiter Glo Assay (Promega, USA). In brief, cells were incubated with or without membrane dye (DiO, 1 uM; Thermofisher Scientific, Australia) for 15 minutes before centrifugation (1200 rpm, 3 minutes) and resuspension in DMEM + 10% FBS. Cells were then seeded at 2 x 10^3^ cells / 200 uL / well, (8 x 10^2^ cells for H460) in ultra-low adherent round-bottom 96-well plates (Lonza, Australia) for a total of 72 hours. Doxorubicin (20 - 50 uM) was added 24 hours post seeding, and silica nanoparticles (SiNP; 10, 30, 100 nm) added (100 ug / mL) in triplicate to spheroids with or without DiO, 48 hours post seeding. At 72 hours, spheroids were transferred in 50 uL total volume to white flat-bottom 96-well plates in triplicate. Media controls were used per each condition in duplicate. An ATP standard curve was also pipetted in duplicate using 10 mM ATP (Life Technologies, Australia) diluted in DMEM + 10% FBS to a range of 0.3125 uM – 10 uM. CellTiter Glo reagent (Promega) was added at 1:1 ratio and plates transferred to an Orbit 4 Benchtop shaker (60 rpm, 30 minutes). ATP concentrations were then measured using 1.0 second luminescence exposure on a Wallac3 Victor plate reader (PerkinElmer, USA). For the live/dead assay, U87, SK-N-BE(2) or H460 cells were seeded at densities mentioned above in ultra-low adherent round-bottom 96-well plates (Lonza) for 72 hours. For EtOH treated controls, 100 uL of 70% EtOH was added to spheroids 1 hour prior to embedding. Spheroids were then washed twice with PBS to remove esterase activity from residual FBS. Spheroids were then gently embedded in sterile, molten 1% low-melt agarose (Sigma) in glass-bottom 24-well plates (Cellvis LLC). These were then treated with 10 µM Ethidium Homodimer-1 and 5 uM Calcein AM from a Live/Dead Test Kit (Molecular Probes, Thermofisher Scientific) to a total volume of 250 uL PBS and incubated (37°C, 95% humidity and 5% CO_2_) for 30 minutes. Spheroids were then imaged on a Zeiss Celldiscoverer 7 with a 5X/0.35 Plan-Apochromat objective and 2X Tubelens Optovar. Images were acquired with 0.457 µm to pixel scale in XY and 2.8 µm to pixel scale in Z, in three channels sequentially with 394/490/573/691 beam splitters (Channel 1: LED-module 470 nm, 6.45%, 134 ms exposure, emission 514 LP; Channel 2: LED-module 567 nm, 50.05%, 500 ms exposure, emission 617 LP; Channel 3: Brightfield TL LED Lamp, 0.10%, 4 ms exposure). Images were exported to Zen Black (2.1 SP3, Zeiss), processed to normalize channel intensities relative to EtOH treated controls and then exported as TIFF files.

### Confocal microscopy: spheroid preparation and image acquisition

For live confocal experiments, cells were pre-stained with DiO (Sigma Aldrich) (1 uM, 15 minutes, 1200 rpm, 3 minutes) or left unstained. These cells were then seeded at densities as above in ultra-low adherent round-bottom 96-well plates (Lonza) for 60 hours. Spheroids (with or without DiO) were then gently embedded in sterile, molten 1% low-melt agarose (ThermoFisher Scientific) in glass-bottom 24-well plates (Cellvis LLC). Phenol red free media + 10% FBS was added to all wells for imaging. For nanoparticle uptake, nanoparticles were diluted in phenol red free media to 40 ug / mL. These solutions were then added to wells with spheroids (stained with or without DiO) at a 1:1 dilution to a final concentration of 20 ug / mL in 500 uL. PBS (500 uL) was added to outer wells to reduce evaporation and drift during imaging. Additional untreated spheroids (with or without DiO) were used as controls.

Imaging was conducted in Zen Black 2.3 SP1 (Zeiss, Germany) on a Zeiss LSM 880 inverted laser scanning confocal microscope equipped with a FAST Airy scan detector and incubation (37°C, 5% CO_2_). Acquisition was carried out using a Plan-Apochromat 10x/0.45 M27 objective, zoom of 1.5 to 1.7 times, maximum scan speed, and pixel arrays of 1292 by 1292 to 1528 by 1528. Acquisition setup was maximized for resolution but prioritized for time (30-minute acquisition window) to capture temporal fluorescence using frame-fast Airy with two channels simultaneously (Channel 1: 488 nm, 15% laser; Channel 2: 561 nm, 20% laser), 488/561/633 beam splitters and 495-550 BP/ 570 LP filters. Spheroid positions were saved, and the acquisition acquired z-stacks (optical section thickness 1.535 µm) from the core to the circumference (one hemisphere, up to 270 µm) every 30 minutes for a total of 24 acquisitions (total 12 hours). Raw data was saved and exported to Zen Black (2.1 SP3, Zeiss, Germany) for 3D Airy processing (automatic strength of 6.0) before post-processing for maximum intensity projections, orthogonal images, and subsequent analyses.

### Azimuth averaging and nanoparticle quantification by diffusion

Analyses (mid-plane, 3D azimuth and diffusion) were conducted using custom scripts in MATLAB (R2020a, MathWorks, Natick, USA) which built upon previous methods [29]. These are included with user friendly comments (%%) in Supplementary files, and are supported by a User Guide, also in the Supplementary.

In brief, midplane radial averaging was calculated from a user defined coordinate for the center of the spheroid. The membrane channel was used to create a binary mask and define the circumference of the spheroid at each time point. Radii were then defined pixel per pixel from the center to the circumference and nanoparticle intensity averaged and exported at 10 µm intervals into an Excel spreadsheet for graphing using GraphPad Prism (V 9.0.1).

For azimuthal averaging of the 3D dataset (approximately a spheroid hemisphere), data was first resized by a factor of three for ease of processing, the XYZ coordinates of the spheroid core defined by the user and a binary mask generated in 3D of the spheroid circumference. This was then used to define azimuth radii, correcting for reduced pixel resolution in Z (approximately three times that of XY). Once radii were defined, nanoparticle intensity data was calculated for a given radius from the spheroid center, and then interpolated to sample intensities over the linear range of radii (at 10 µm intervals). Data was exported into a Microsoft Excel spreadsheet and imported into GraphPad Prism (V 9.0.1) for further processing.

For kinetic quantification, radially averaged and time evolving intensity profiles were used to extract particle diffusivity by the physical principles that are often employed in fluorescence recovery after photobleaching (FRAP) [65]. Briefly, for each radius from the spheroid center, the time evolution of average intensity in particle channel, was fitted with a forward in time, central in space (FTCS) diffusion model in 2D [38].

For each radial distance from the spheroid center, *w*, we extracted this way a diffusion time, *t_D_*, and from these two parameters were able to calculate particle diffusivity using the relationship in equation (1):

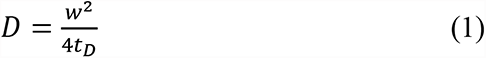

Data outputs were saved in Microsoft Excel spreadsheets and imported into GraphPad Prism for graphical sketching and statistics.

### Mathematical modelling to calculate diffusivity – Model development

To calculate nanoparticle diffusion, we first considered the evolution of the number of nanoparticles, N(r,q,f,t), as a function of radial distance from the center of the tumor spheroid, r, the polar angle, q, the azimuthal angle, f and time, t. We made the assumption that the tumor spheroid can be approximated with a sphere, and that the number of nanoparticles does not depend on the orientation of the sphere. As such, we ignored the azimuthal and polar angles, and hence the number of nanoparticles, N(r,t) depends only on radial distance and time. We assumed that there was direct proportionality between the number of nanoparticles and relative, measured, fluorescence and given that all the imaging parameters were kept consistent, we could ignore the constant terms in the solution of the problem.

In this model we derive nanoparticle motion as primarily driven by diffusion. It is likely that nanoparticle motion is affected by the cells within the spheroid, and that cells may behave differently depending on distance from the center of the spheroid, as oxygen levels can decrease toward the center of the spheroid. Accordingly, we allow the diffusion function to vary as a function of the radial distance. This could correspond to changes in cell density or cell behavior as a function of the radial distance.

However, we do not specifically state the biological behavior behind potential changes in nanoparticle diffusion, we merely assume that diffusion can vary as a function of the radial distance. As such, the evolution of the number of nanoparticles is described by the following partial differential equation (PDE) (2);

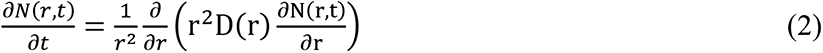

where *D(r)* is the diffusivity of the nanoparticle as a function of radial distance. Outside of the spheroid, that is*, r > r_sph_*, the nanoparticle diffusivity will be equal to the diffusivity given the Stokes-Einstein equation, *D_SE_*, in equation (3),

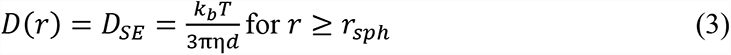

where *k_b_* is the Boltzmann constant, *T* is the temperature, *h* is the dynamic viscosity and *d* is the diameter of the nanoparticle. We assumed that the diffusivity of the nanoparticle in the spheroid is reduced compared to the Stokes-Einstein equation. Further, we assumed that the nanoparticle diffusivity is most inhibited toward the center of the spheroid, where cell function may be impacted most significantly. We therefore make the choice that the nanoparticle diffusivity is described by equation (4),

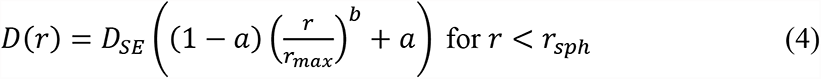

where the parameter *a* and *b* are determined by the data. This choice of diffusivity function allows for a monotonic increase in diffusivity with radial distance, consistent with our assumptions, with the minimum diffusivity at the center of the spheroid given by *D(0) = aD_SE_*. The rate of increase of diffusivity is controlled by the parameter *b*, where *b* = 1 corresponds to a linear increase in diffusivity, for example. Importantly, at *r = r_sph_*, the nanoparticle diffusivity is equal to the diffusivity given by the Stokes-Einstein equation. This choice of diffusivity function allows for flexibility, while still incorporating known nanoparticle behavior and minimizing the number of free parameters to be determined from the data. At the boundaries of the domain, at *r =* 0 and *r = r_max_* we made the assumption that, on average, a nanoparticle is equally likely to enter the domain as it is to leave the domain. This corresponded to a zero-flux boundary condition. At the beginning of the simulation, *t* = 0, we set the number of nanoparticles inside the spheroid to be zero, with a constant number of nanoparticles outside of the spheroid, consistent with the experimental conditions.

### Mathematical modelling to calculate diffusivity – Solution method

To obtain a solution to the PDE (2) governing the number of nanoparticles we first spatially discretized the governing PDE onto a uniform grid with spacing *Dr* via a central difference approximation for the spatial derivatives. We defined this grid between *r* = 0 and *r* = *r_max_* = *Kr_sph_* where *K* was chosen such that the boundary of the solution domain is sufficiently far away from the boundary of the spheroid and *r_sph_* = 250 mm. For all experimental datasets we chose *K* = 7 and we verified that the solution is not sensitive to increases in K. We selected a backward Euler approximation with constant timesteps of length *Dt* to approximate the temporal derivative. We solved the PDE (2) up to *t* = 12 h, consistent with the experiment, and select *Dt* such that we had 2400 time steps, *i.e. Dt* = 0.005 h. We solved the resulting system of tridiagonal equations using the Thomas algorithm.

### Mathematical modelling to calculate diffusivity – Parameter estimation

To determine experiment-specific values of *a* and *b*, we fitted the numerical solution of the PDE (2) to the experimental data. Due to the three-dimensional averaging process, there can be a drop in fluorescence for sufficiently large *r* values if the spheroid is not perfectly symmetric and hence, we considered data for *r* values until we observe this drop.

We obtain the numerical solution for particular *a* and *b* values and compare the solution at the *r* values where we have experimental measurements. Using MATLAB’s *lsqnonlin* function, which implements the Levenberg-Marquadt algorithm, we determined values of *a* and *b* that best fit the data for each experiment. Further, we verified that the predicted number of nanoparticles (or, equally, the level arbitrary fluorescence) matched the experimental data well. For each experiment we found that our model described the experimental data well. We determined *a* and *b* values for each replicate of an experiment and report the mean and standard error of *a* and *b* for each nanoparticle-spheroid combination.

### Particle diffusion modelling in simulated spheroids

To validate our analysis (DONUTS), we simulated spheroids of varying cell densities and sizes where particles were simulated to move from outside towards the core of spheroids. Simulated spheroid cellular positions in 3D were generated using the DistMesh package [66] and subsequent particle motion was overlayed over time in MATLAB using custom scripts available in Supplementary (R2020a, Mathworks, Natick, USA). The simulator allowed for varying cell and spheroid size as well as particle density, their diffusivity inside and outside of cell objects and the probabilities for particles to move in and out of cell objects. The simulated volume size was set to 300 pixels in X, Y and Z, number of timepoints was set to 1000 frames, pixel size to 0.1 µm and frame time to 1 s. To simulate the asymmetry in point spread function (PSF), the full width half maximum (FWHM) in XY was set to half that of the axial PSF FWHM (in Z). Within the total image volume, the spheroid volume was set to 50% of the total image space, enabling necessary free space for particle generation and directional uptake into the spheroid. Cell density fraction (CDF) defined the occurrence of the centroid of cell objects relative to the total spheroid, while cell radius (*r*) defined the total cell size relative to these centroids, maintaining that CDF ≥ 2*r* to prevent object overlap. CDF was varied to alter object density, while *r* was altered to cell object size, generating three convolved spheroids with CDF and *r* defined in Table **1**.

**Table 1.**
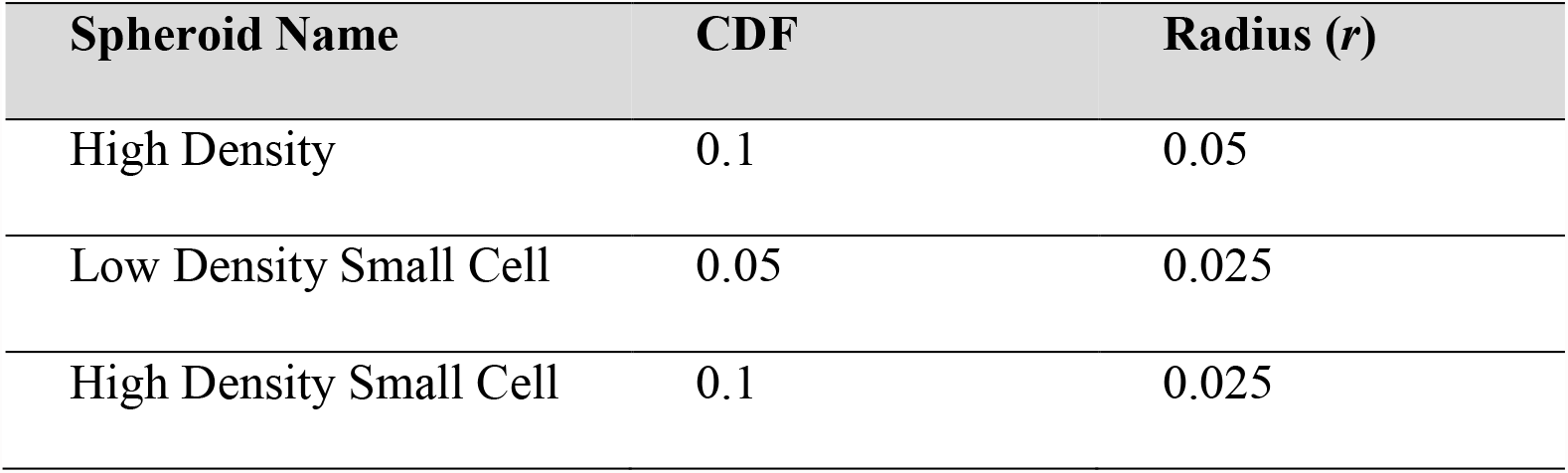
Cell density fraction (CDF) and radius of convolved (simulated) spheroids used to model particle diffusion as a method of external validation of DONUTS.

| Spheroid Name | CDF | Radius (*r*) |
| --- | --- | --- |
| High Density | 0.1 | 0.05 |
| Low Density Small Cell | 0.05 | 0.025 |
| High Density Small Cell | 0.1 | 0.025 |

Final parameters assumed set diffusion coefficients for particles inside cells (D(in)) and particles outside or between cells (D(out)) as well as associated probabilities of whether a particle would enter a cell object, and if so, the subsequent probability of exiting cell objects. D(in) was assumed to encompass all intracellular mechanisms of particle uptake and trafficking and was the only particle-related variable altered during simulations. Scripts are available in Supplementary.

### Lightsheet microscopy: cell preparation and image acquisition

Live spheroids of glioblastoma (U87), neuroblastoma (SK-N-BE(2)) and NSCLC (H460) were prepared as above. After 72 hours, spheroids were transferred into 150 uL aliquots of fixing solution (4% paraformaldehyde (ProSci Tech PtyLtd, Australia), 0.1% glutaraldehyde (Sigma) in 1% PBS) and fixed for 5 – 7 days (4°C, gentle agitation at 70 rpm). Fixed spheroids were then embedded in 2% low melt agarose (Sigma) using a hollow plastic cylinder from FEP HS 0.125 EXP/0.086 REC Fusing Sleeve (Zeus Virtual Sample Locker) as previously described [67]. Agarose embedded spheroids were then transferred into Cubic L solution (10% Triton X-100, 10% N-buthyldiethanolamine in MilliQ water (w.w), Sigma) for 3 – 4 days (37°C, 60 rpm agitation). Samples were washed thrice (PBS, 2 hours) before immersion in 1% PBS containing 1 µM DAPI for 24 hours (37°C, 60 rpm). Samples were washed thrice in PBS as above and finally transferred into Cubic R solution (45% antipyrine, 30% nicotinamide in MilliQ water (w.w), Sigma) for a minimum of four days prior to imaging. Lightsheet imaging was conducted in Zen Black 2014 SP1 (Zeiss) on a Zeiss Lightsheet Z.1 with 1.45 N cubic corrected Plan Neofluar 20X/1.0 objective and 10X/0.2 LSFM clearing lateral objectives. Magnification was adjusted to 1.0 times with a 1920 by 1920-pixel grid. Z-stack step size was set to optimal (0.390 – 0.412 µm) and images acquired with a 405 nm laser (2.0%, 99.95 ms exposure), 405/488/561/640 laser block filter and emission 460-500 BP. Images were exported for processing in Zen Black (2.1 SP3, Zeiss), with Dual Side Fusion of left and right lasers using Maximum Intensity Fusion. Analysis for single nuclei detection was done as referenced in [41] and further quantification of Nearest Neighbor Distances (NND) between nuclei was done in custom built scripts in MATLAB (2020a, MathWorks, Natick, USA). Script is available in Supplementary, and full instructions for reproduction and use available in User Guide, also in the Supplementary Material.

### Force imaging cytometry for cell diameter, deformation and stiffness

The diameter, stiffness and deformation of U87, SK-N-BE(2) and H460 single cell suspensions was measured using force imaging cytometry and real-time deformation, as described previously [68]. Briefly, cells were harvested from a 70-90% confluent flask as described in culture conditions above, counted using Trypan Blue and resuspended in 1-2 mL PBS at 3 x 10^5^ cells / mL. These cells were then centrifuged as above and gently resuspended in CellCarrier A buffer (Zellmechanik Dresden) before being transferred into a Falcon^TM^ round-bottom polystyrene test tube with cell strainer cap (Corning) to ensure single cell suspension. Samples were loaded onto a syringe pump (neMESYS; Cetoni), AcCellerator L1 system (Zellmechanik Dresden) with synchronized pulsed LED illumination. This system was built into a Zeiss AxioObserver (Zeiss, Germany) with 40X/0.65 objective, CMOS camera, with a 1024 x 1280 pixel grid, 8 bit imaging depth, maximal resolution of 340 nm per pixel and frame rate of 4000 frames s^−1^. Samples were run through 30 µm microfluidic chips at a cell flow rate of 0.0400 µL s^−1^ and sheath flow rate of 0.120 µL s^−1^ using Shape In (2.2.2.4). Hard area gates were set at 50 – 200 µm^2^ depending on cell size to exclude particulate matter. Raw data was exported and analyzed in Shape Out (2.6.4), with manual curation to gate true single cell populations, followed by calculation of Young’s Modulus, deformation according to equation (5) [68].

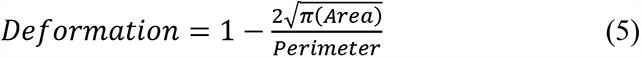

Statistical analysis was performed in PRISM (9.0.1).

### Data Statement

All data presented in this manuscript is available from the corresponding author on request. All custom scripts have been made available at [GitHub link prior to publication], along with the User Guide for installation of MATLAB and walkthrough use of the DONUTS analysis package. Supporting analysis packages in MATLAB are also included. These can be accessed via GitHub above, or in the zipped folder in the Supplementary of this paper.

#### Supporting Information Available

Supplementary data (figures and tables) are included which support data included in the primary manuscript. In addition, an analysis package for DONUTS is included as a .zip file. This contains the code for DONUTS analysis scripts for use in MATLAB and a READ ME User Guide which provides step-by-step instructions for the installation and use of all analysis packages detailed in the manuscript, with the intention to assist general use of this data analysis platform. Embedded in the User Guide are also four sample datasets accessible through FigShare, which can be used to test and validate our analysis platform. This guide also contains information to assist in initial experimental setup and trouble-shooting recommendations.

## AUTHOR CONTRIBUTIONS

MK conceived the initial project and provided funding and supervision, with TPD and JM. MK, EJC and RMW provided experimental, modelling and imaging resources, respectively. AA-C refined the project conception, developed methodologies, and performed all experiments, with support from EP, CH, FMM and RMW. AA-C and EP conceived the analysis method DONUTS and EP developed analysis scripts. AA-C and EP performed validation of DONUTS analysis. STJ developed applied mathematical modelling analysis and investigated nanoparticle penetration kinetics, with support from EJC. AA-C prepared the figures and wrote the manuscript. FMM, RMW and MK reviewed and edited versions of the manuscript. EP and STJ reviewed the manuscript and provided additional input for methodology and analysis. All authors read and approved the final manuscript.

## ACKNOWLEDGMENTS

This work was supported by the Children’s Cancer Institute, which is affiliated with the University of New South Wales (UNSW Sydney), and the Sydney Children’s Hospital Network, and by grants from the National Health and Medical Research (Program Grant APP1091261 and Principal Research Fellowship APP1119152 to MK) and Cancer Institute New South Wales Program Grant (TPG2037 to MK). MK is also supported by Australian Research Council Center of Excellence in Convergent Bio-Nano Science and Technology (CE140100036). AA-C acknowledges support from the Scientia PhD Scholarship Scheme (UNSW Sydney), the Josee Hilton Excellence Award and Children’s Cancer Institute Postgraduate Top-up Scholarship. STJ is supported by the Australian Research Council (DE200100988). The authors would like to thank the Katharina Gaus Light Microscopy, at the Mark Wainwright Analytical Center for their support and resources involved in this work, and the Cancer Institute of NSW for a donation that allowed the acquisition of a Zeiss LSM 880 used in this work. The authors would also like to thank J. Fletcher for his mnemonic which refined the name of this analysis method.

\# Special acknowledgement of our colleague and dear friend, Professor Edmund Crampin, who passed away during the writing of this manuscript.

## SUPPLEMENTARY DATA

### Supplementary Figures

**Supplementary Figure 1:**
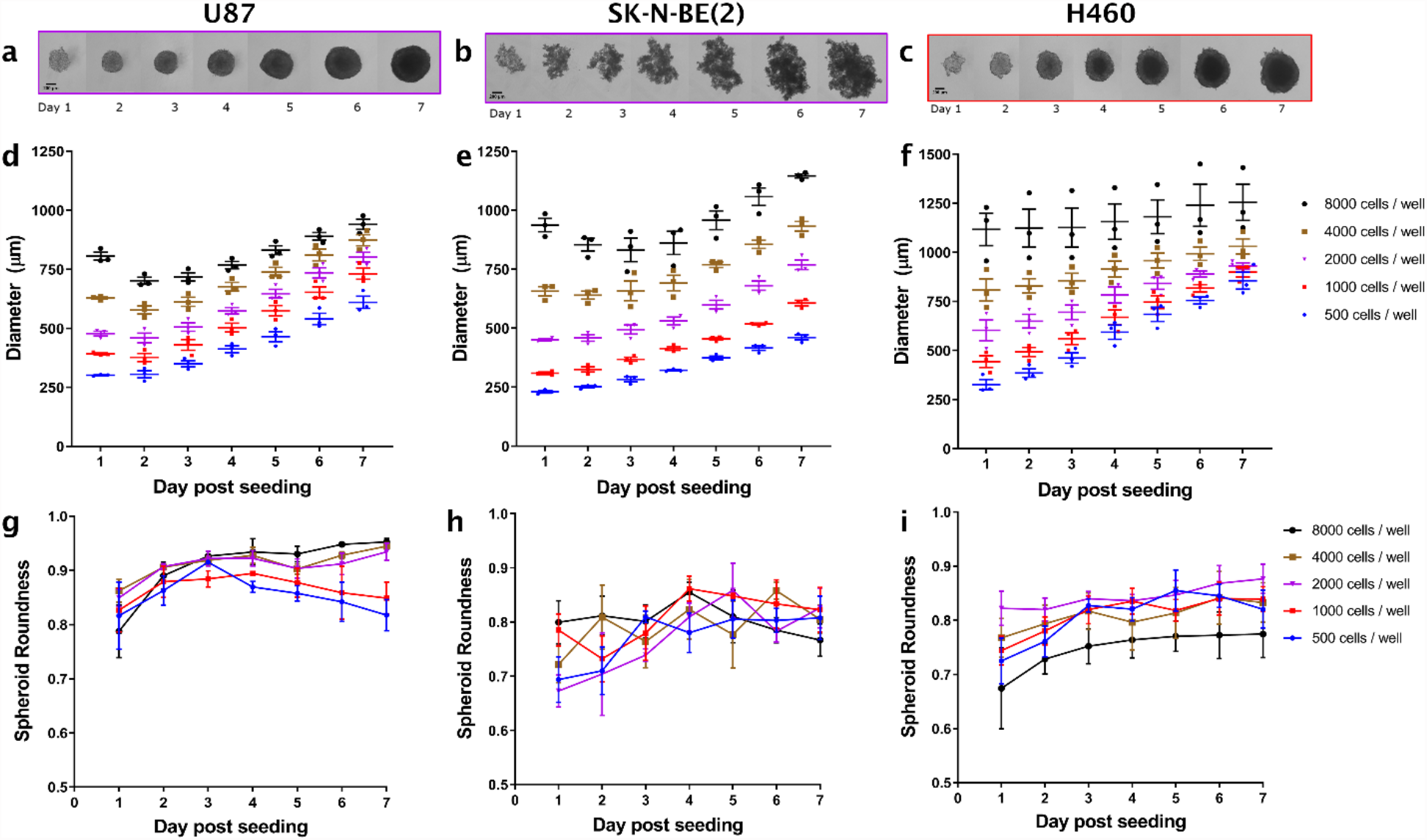
Spheroid growth of glioblastoma (U87), neuroblastoma (SK-N-BE(2)) and non-small cell lung cancer (H460) seeded at Day 0 and imaged daily for 7 days. **(a)** Representative growth of U87, **(b)** SK-N-BE(2) spheroid (2000 cells at Day 0). **(c)** Representative growth of H460 spheroid (1000 cells at Day 0). Scale bar, 200 µm. This growth was quantified in ImageJ by measuring the diameter each day in **(d)** U87, **(e)** SK-N-BE(2) and **(f)** H460 spheroids at different seeding densities indicated in (f). Points, individual biological replicates (n=3) per time point, per seeding density. Lines, mean of n=3 ± SEM. Growth characteristics were quantified by aspect ratio, defined above as roundness in **(g)** U87, **(h)** SK-N-BE(2) and **(i)** H460 spheroids at the same seeding densities as above. For ease of visualisation, points here represent mean of n=3. Bars, SEM.

**Supplementary Figure 2:**
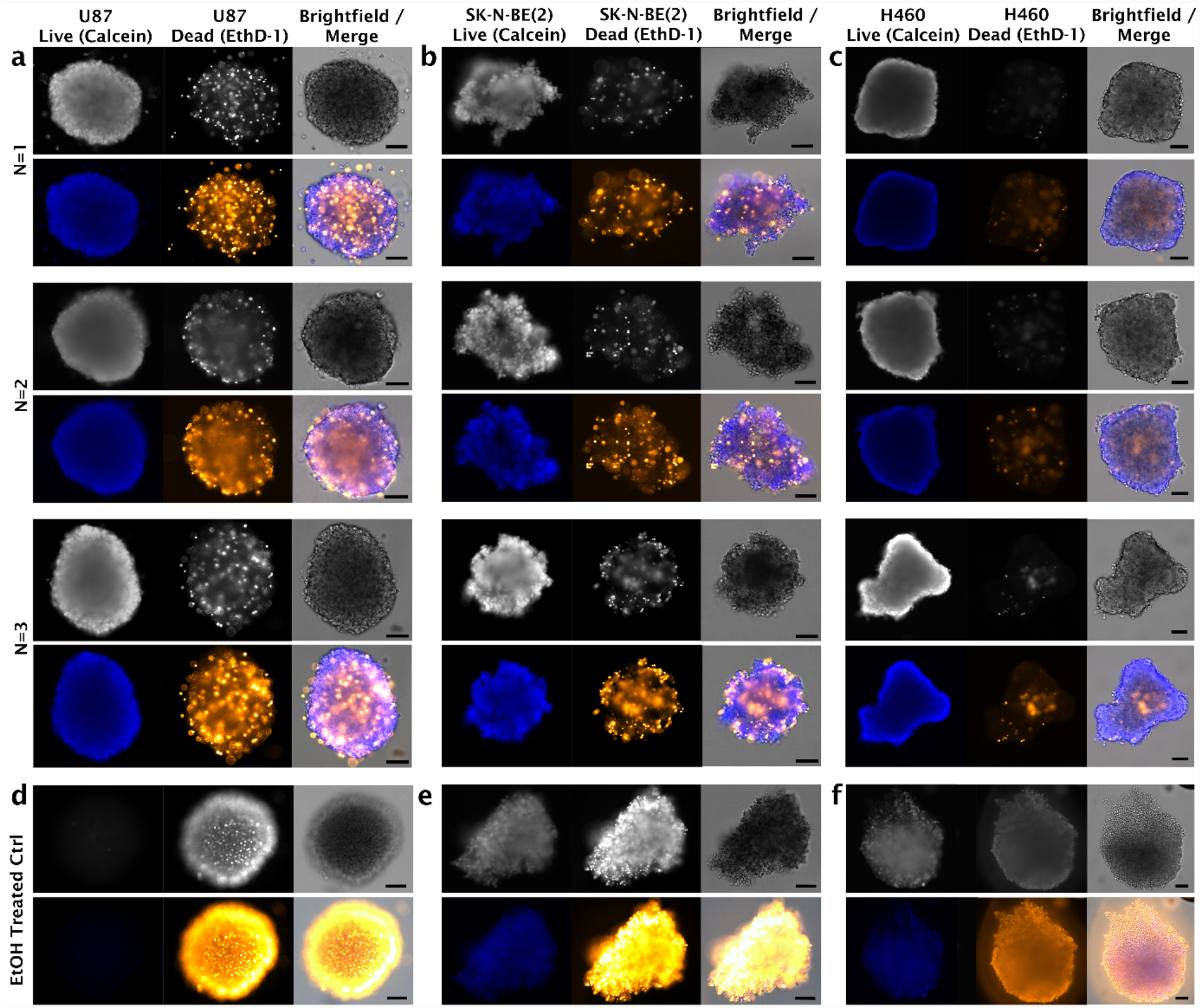
Spheroid viability and characteristics visualized through a live/dead assay using Calcein and ethidium homodimer-1 (EthD-1). Representative brightfield and live (blue)/dead (orange-yellow) images of **(a)** U87 and **(b)** SK-N-BE(2) and **(c)** H460 cell spheroids, which were grown for 3 days in low adherent round-bottom well plates with an initial seeding density of 2 × 10^3^, or 8 x 10^2^ cells for H460 specifically. Viability is contrasted against ethanol (EtOH) treated controls for **(d)** U87 and **(e)** SK-N-BE(2) and **(f)** H460 respectively. Scale bar, 100 µm.

**Supplementary Figure 3:**
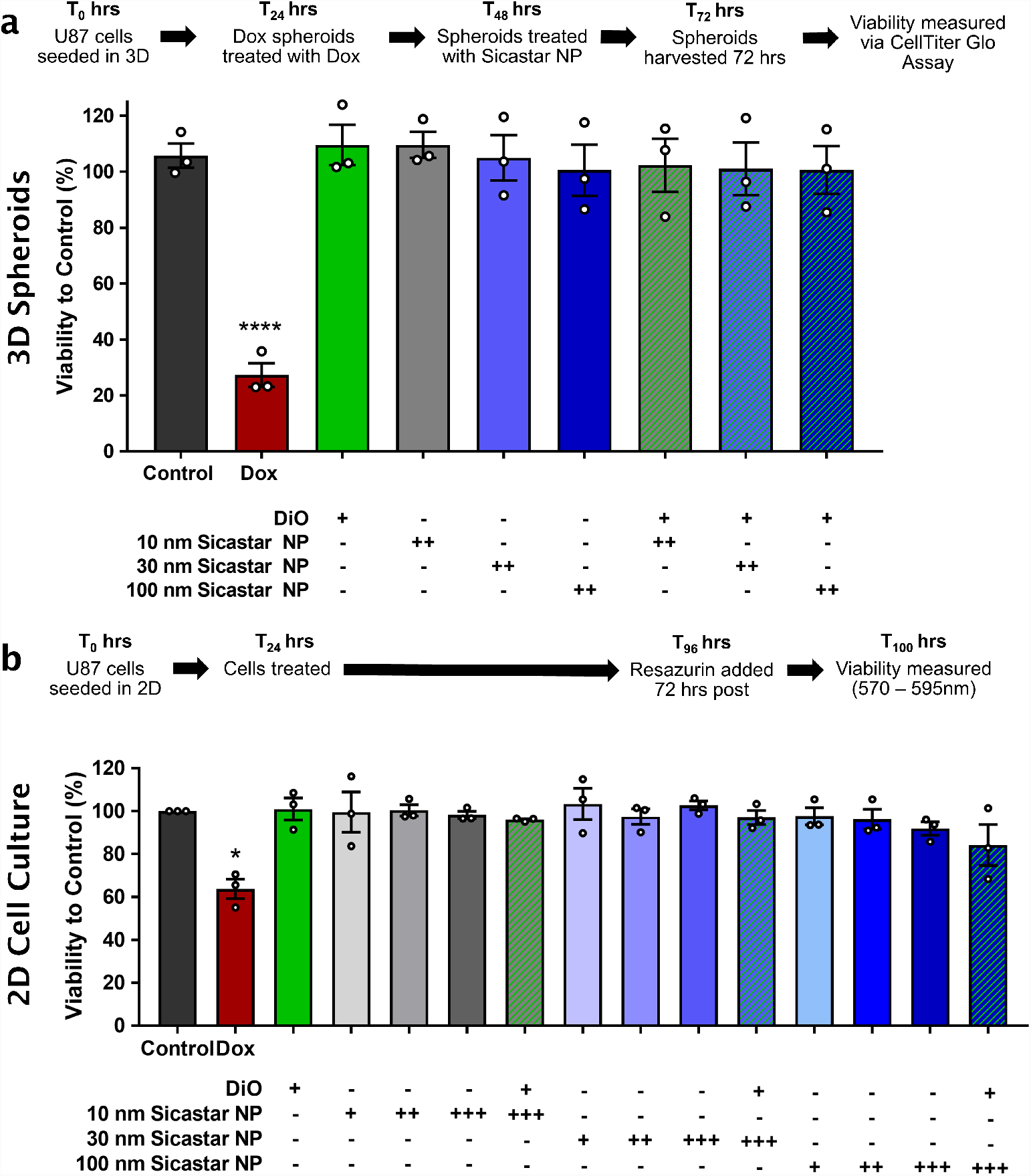
Cell viability of glioblastoma (U87) cells treated with silica nanoparticles (SiNPs) and membrane dye (DiO). **(a)** 3D cell viability of U87 following treatment SiNPs alone or in combination with membrane dye (1 µM) for 24 hrs. Measured using Celltiter Glo Assay. Doxorubicin (Dox, 50 uM) used as a positive control. *Points*, biological replicates. *Columns,* mean of n = 3. *Bars,* SEM. Significance to control (untreated) using one-way ANOVA, **** *p* < 0.0001. **(b)** 2D cell viability of U87 following treatment with SiNP (+ 1 ug / mL, ++ 10 ug / mL, +++ 100 ug / mL, as indicated) membrane dye (1 µM) and combination over 72 hours. Doxorubicin (Dox, 0.25 uM) used as a positive control. Measured using a Resazurin-based cell viability assay. *Points*, biological replicates. *Columns,* mean of n = 3. *Bars,* SEM. Significance to control (untreated) using a paired ratio t-test, * *p* < 0.05.

**Supplementary Figure 4:**
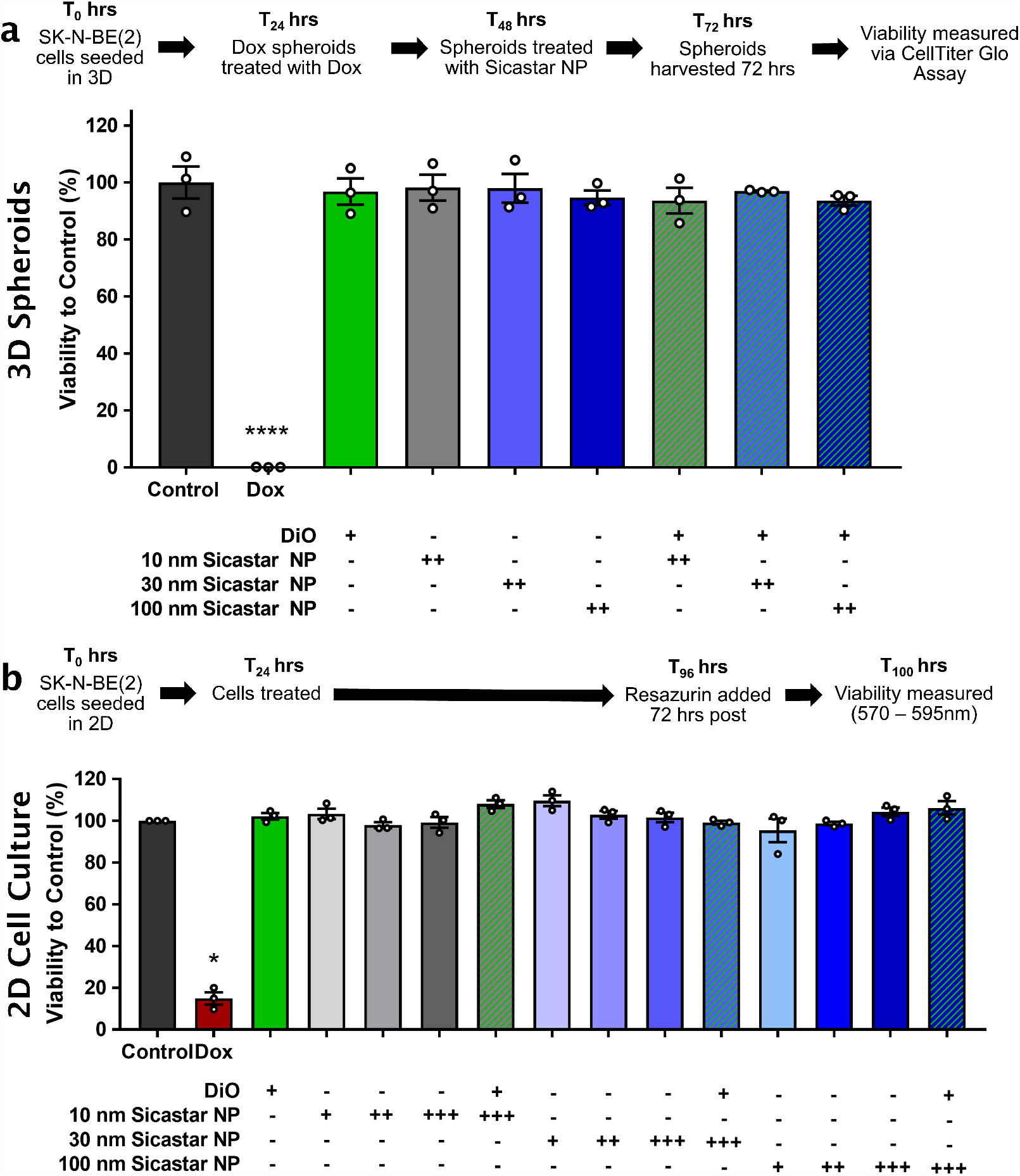
Cell viability of neuroblastoma (SK-N-BE(2)) cells treated with silica nanoparticles (SiNPs) and membrane dye (DiO). **(a)** 3D cell viability of SK-N-BE(2) following treatment SiNPs alone or in combination with membrane dye (1 µM) for 24 hrs. Measured using Celltiter Glo Assay. Doxorubicin (Dox, 50 uM) used as a positive control. *Points*, biological replicates. *Columns,* mean of n = 3. *Bars,* SEM. Significance to control (untreated) using one-way ANOVA, **** *p* < 0.0001. **(b)** 2D cell viability of SK-N-BE(2) following treatment with SiNP (+ 1 ug / mL, ++ 10 ug / mL, +++ 100 ug / mL, as indicated) membrane dye (1 µM) and combination over 72 hours. Doxorubicin (Dox, 0.25 uM) used as a positive control Measured using a Resazurin-based cell viability assay. *Points*, biological replicates. *Columns,* mean of n = 3. *Bars,* SEM. Significance to control (untreated) using a paired ratio t-test, * *p* < 0.05.

**Supplementary Figure 5:**
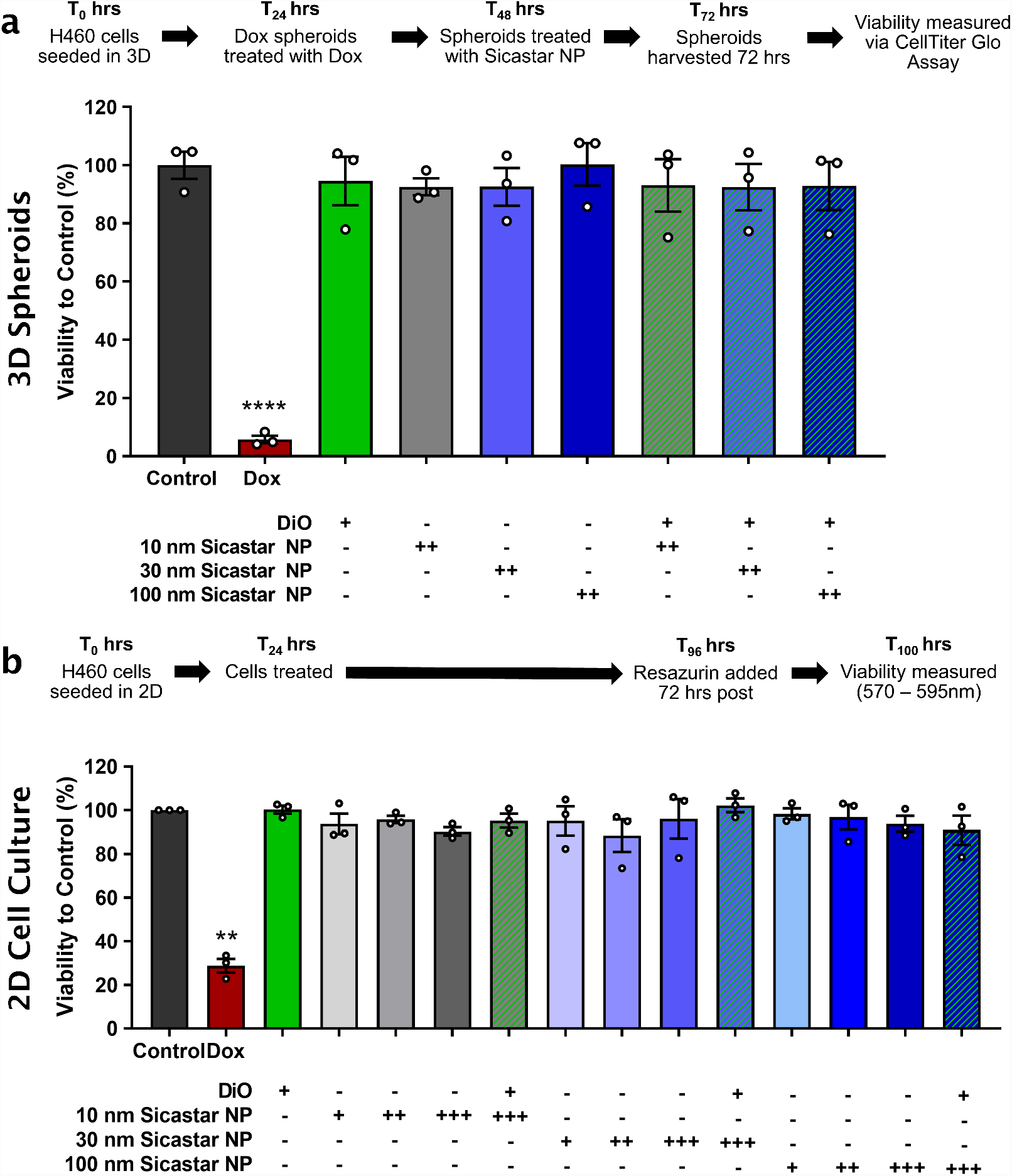
Cell viability of non-small cell lung cancer (H460) cells treated with silica nanoparticles (SiNPs) and membrane dye (DiO). **(a)** 3D cell viability of H460 following treatment SiNPs alone or in combination with membrane dye (1 µM) for 24 hrs. Measured using Celltiter Glo Assay. Doxorubicin (Dox, 20 uM) used as a positive control. *Points*, biological replicates. *Columns,* mean of n = 3. *Bars,* SEM. Significance to control (untreated) using one-way ANOVA, **** *p* < 0.0001. **(b)** 2D cell viability of H460 following treatment with SiNP (+ 1 ug / mL, ++ 10 ug / mL, +++ 100 ug / mL, as indicated) membrane dye (1 µM) and combination over 72 hours. Doxorubicin (Dox, 0.1 uM) used as a positive control. Measured using a Resazurin-based cell viability assay. *Points*, biological replicates. *Columns,* mean of n = 3. *Bars,* SEM. Significance to control (untreated) using a paired ratio t-test, ** *p* < 0.01.

**Supplementary Figure 6:**
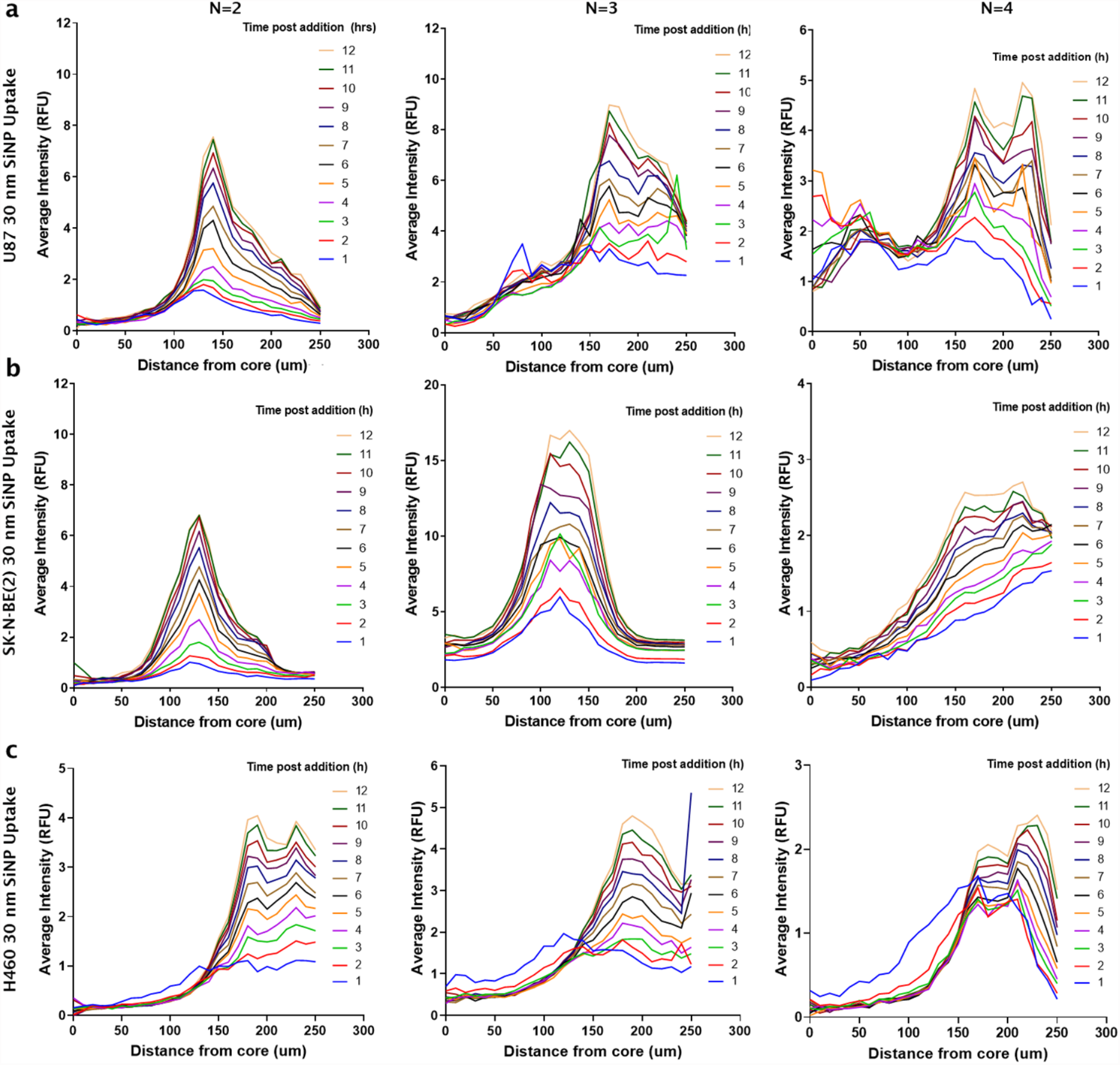
**3D Azimuthal quantification of 30 nm SiNP uptake** in **(a)** glioblastoma (U87), **(b)** neuroblastoma (SK-N-BE(2)) and **(c)** non-small cell lung cancer (H460) cell spheroids. Each graph represents 30 nm SiNP tumor spheroid uptake of a biologically independent experiment, in addition to the data presented elsewhere.

**Supplementary Figure 7:**
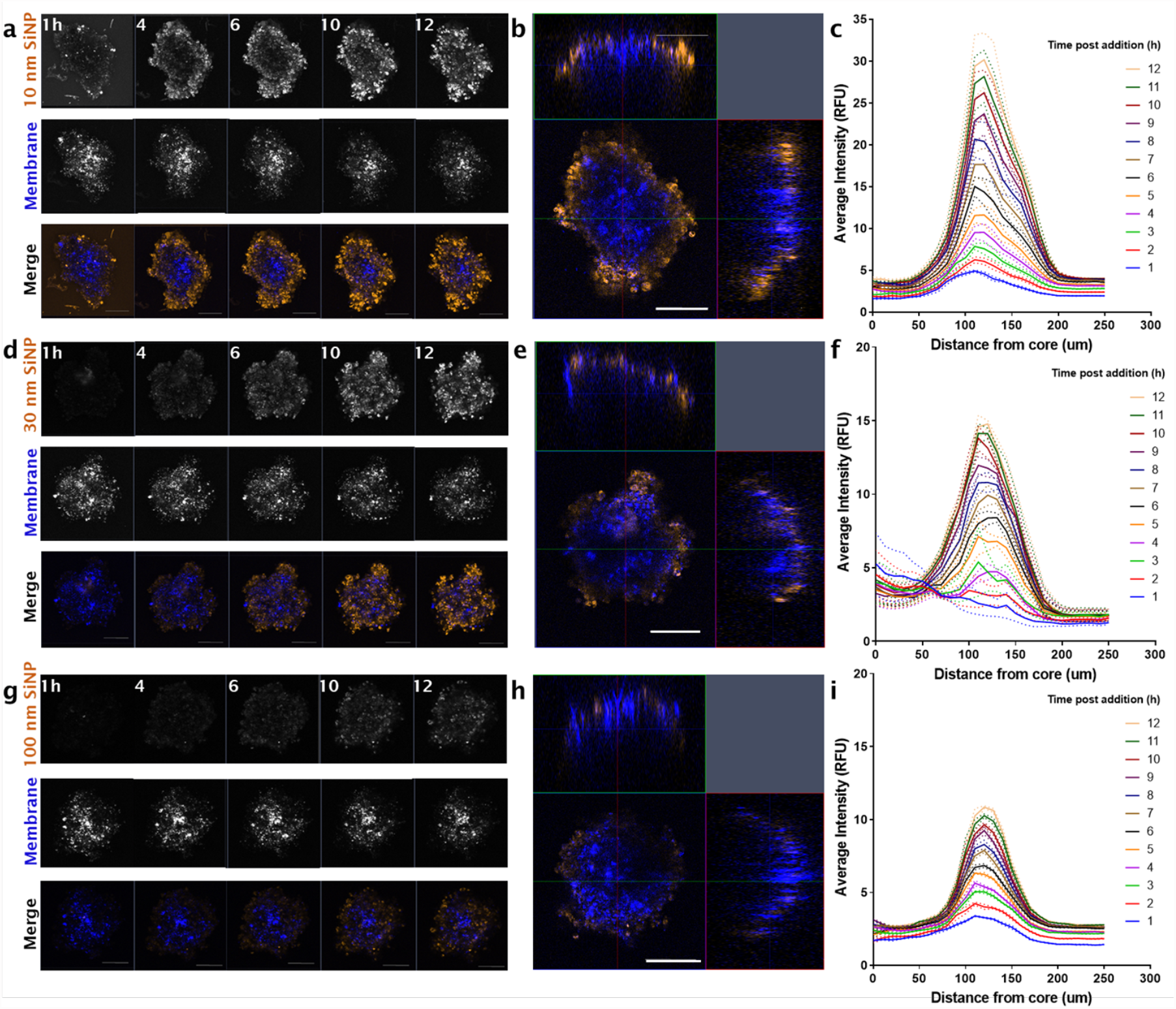
Silica nanoparticle (SiNP) uptake in neuroblastoma (SK-N-BE(2)) tumor spheroids over 12 hours. **(a)** Representative maximum intensity projections of 10 nm SiNP, membrane (DiO) and merge at 1, 4, 6, 10, 12 hours post addition (Orange, SiNP; Blue, Membrane). Z-stack images acquired using a Zeiss 880 confocal microscope (Fast Airy, sequential frame-fast laser excitation at 488 nm and 561 nm, 10X objective). **(b)** Orthogonal (XY, XZ, ZY) merge of SK-N-BE(2) spheroid at six hours post SiNP addition, representative of n = 4. **(c)** Representative quantification of nanoparticle uptake from the core of the spheroid to the circumference over time with increased 10 nm SiNP penetration. Analysis conducted using a 3D azimuth averaging custom script, MATLAB (2020a). Workflow above was performed for 30 nm SiNP showing **(d)** maximum intensity projections over 1, 4, 6, 10, 12 hours post SiNP addition; **(e)** orthogonal merge at six hours and **(f)** azimuth quantification respectively. Imaging and analysis also performed for 100 nm SiNP in panels **(g)** maximum intensity projections; **(h)** orthogonal merge at six hours and **(i)** quantification of 100 nm SiNP uptake. Lines, mean of t=3 analysis iterations. Dotted lines, error SEM. Scale bar, 100 µm.

**Supplementary Figure 8:**
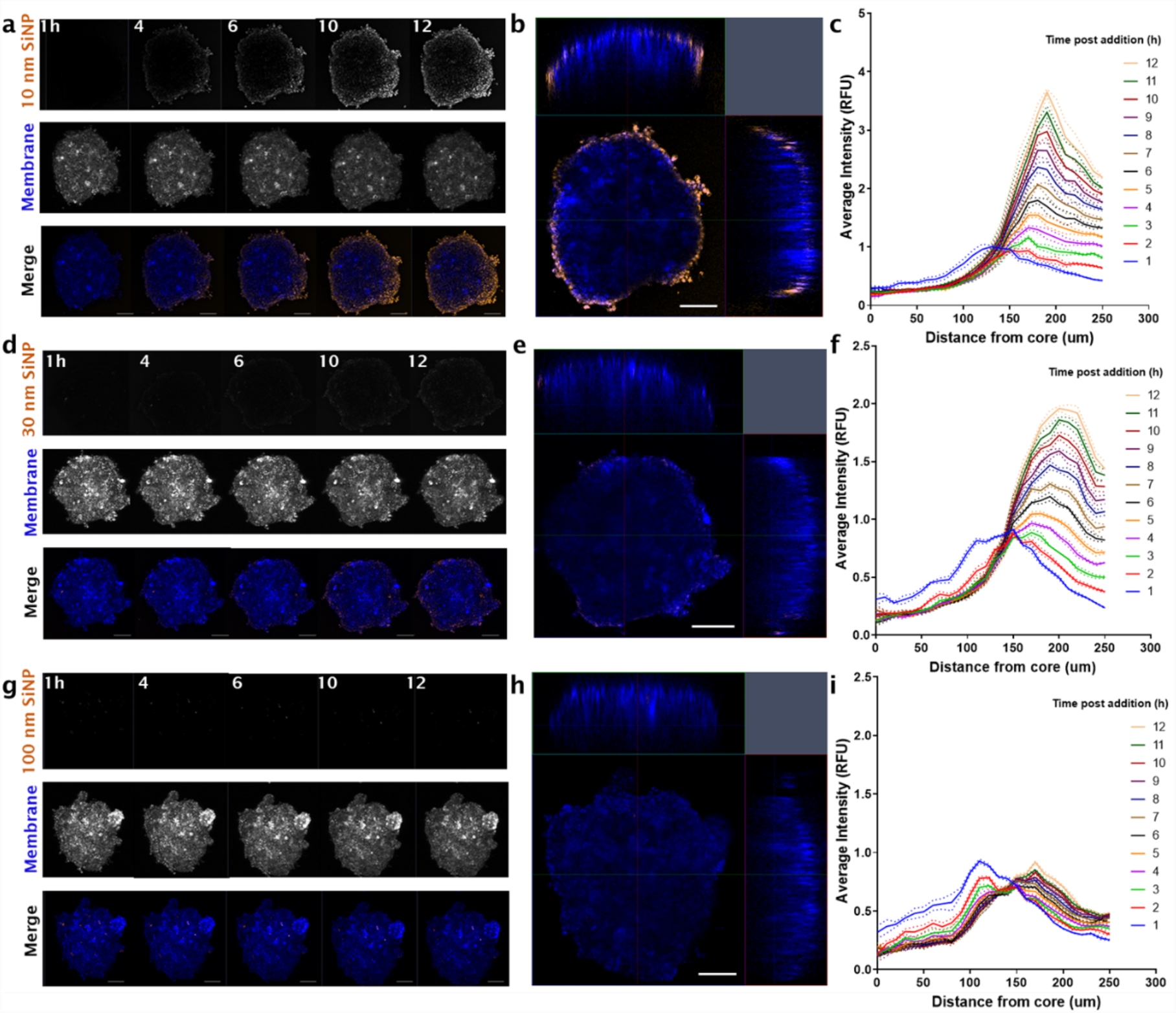
Silica nanoparticle (SiNP) uptake in non-small cell lung cancer (H460) tumor spheroids over 12 hours. **(a)** Representative maximum intensity projections of 10 nm SiNP, membrane (DiO) and merge at 1, 4, 6, 10, 12 hours post addition (Orange, SiNP; Blue, Membrane). Z-stack images acquired using a Zeiss 880 confocal microscope (Fast Airy, sequential frame-fast laser excitation at 488 nm and 561 nm, 10X objective). **(b)** Orthogonal (XY, XZ, ZY) merge of H460 spheroid at six hours post SiNP addition, representative of n = 4. **(c)** Representative quantification of nanoparticle uptake from the core of the spheroid to the circumference over time with increased 10 nm SiNP penetration. Analysis conducted using a 3D azimuth averaging custom script, MATLAB (2020a). Workflow above was performed for 30 nm SiNP showing **(d)** maximum intensity projections over 1, 4, 6, 10, 12 hours post SiNP addition; **(e)** orthogonal merge at six hours and **(f)** azimuth quantification respectively. Imaging and analysis also performed for 100 nm SiNP in panels **(g)** maximum intensity projections; **(h)** orthogonal merge at six hours and **(i)** quantification of 100 nm SiNP uptake. Lines, mean of t=3 analysis iterations. Dotted lines, error SEM. Scale bar, 100 µm.

**Supplementary Figure 9:**
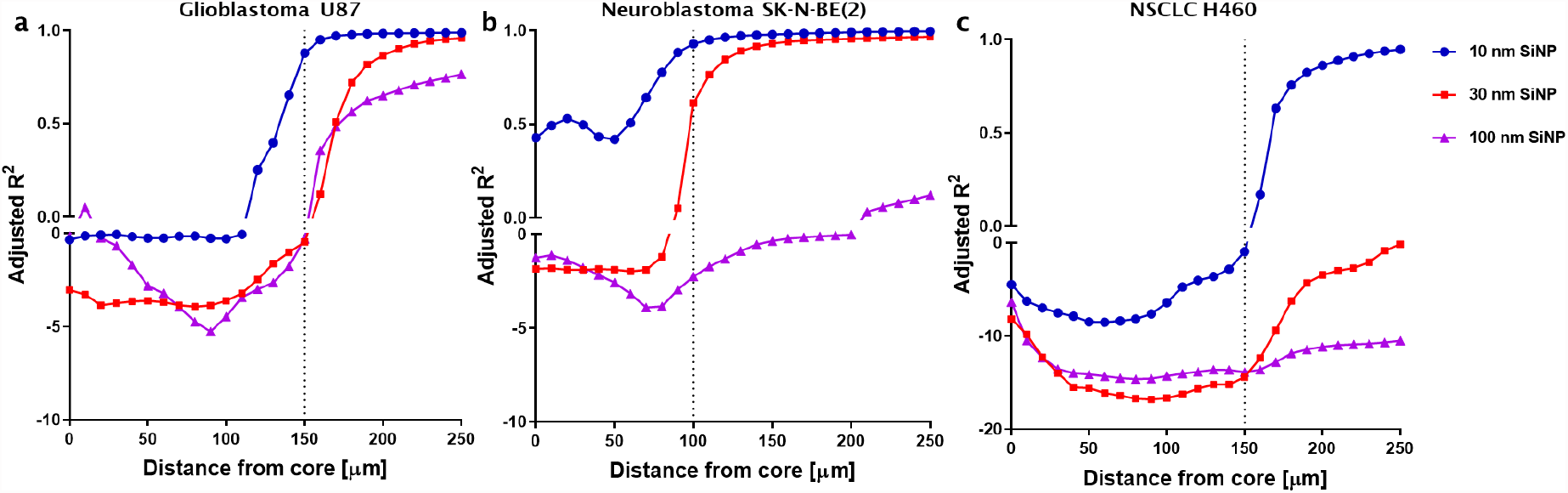
**Adjusted r squared (R^2^) fit values for silica nanoparticle (SiNP) diffusion kinetics** for **(a)** glioblastoma U87 **(b)** neuroblastoma (SK-N-BE(2)) and **(c)** H460 spheroids, calculated using the forward in time, central in space (FTCS) coefficient method. Dotted lines, right side indicating improved fit of model with R^2^ → 0.99, excluding 100 nm SiNP in SK-N-BE(2) and 30 nm, 100 nm in H460 which did not show R^2^ above 0.25.

**Supplementary Table 1:**
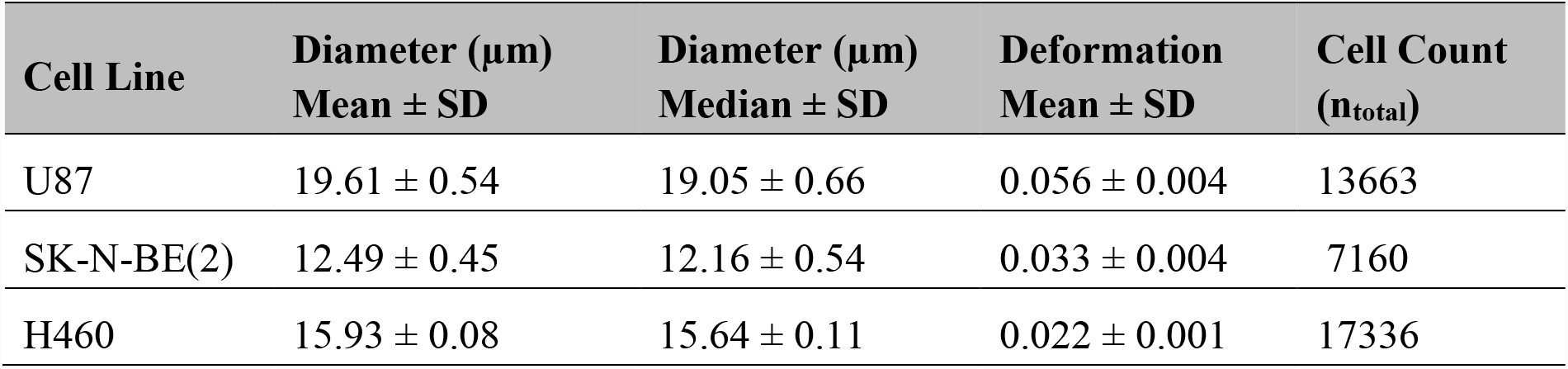
**Cell characteristics of U87, SK-N-BE(2) and H460** with the mean and median diameters, alongside cell counts and deformation. Quantified using single-cell force imaging cytometry at a total flow rate of 0.160 uL s^−1^.

**Supplementary Figure 10:**
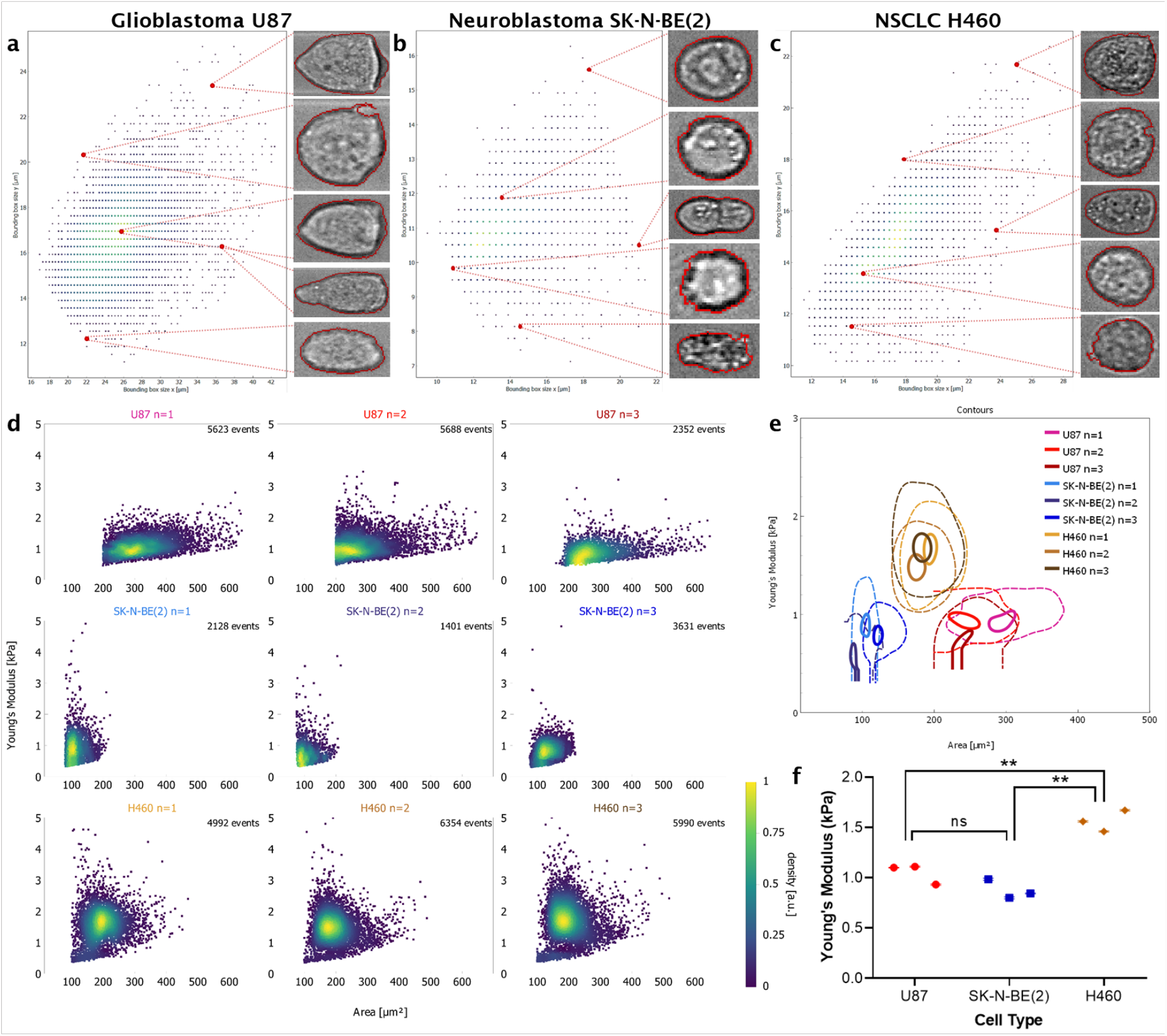
**Single cell stiffness of U87, SK-N-BE(2) and H460 cells,** measured using force imaging cytometry (AcCellerator, Zellmechanik Dresden). Representative images of single cells of **(a)** U87, **(b)** SK-N-BE(2) and **(c)** H460 which were imaged at a total flow rate of 0.16 uL s^−1^. Cells were gated according to size in X and Y as above to ensure single cell suspensions. **(d)** Scatter plots generated of cell area versus Young’s Modulus calculated for each biological run, 1400 minimum cells per run. Heat map representative of count rate across single cell scatter. **(e)** Contour plots of Young’s Modulus for each cell line. *Solid line,* 95^th^ percentile. *Dotted lines*, 50^th^ percentile of total cell population. The average of these values was then used in **(f)** to calculate significant differences of Young’s Modulus between cell types. *Points*, mean of biological replicate. *Bars*, SEM. Significance by unpaired t-tests with Tukey correction, ns, non-significant, ** *p* < 0.01.

**Supplementary Figure 11:**
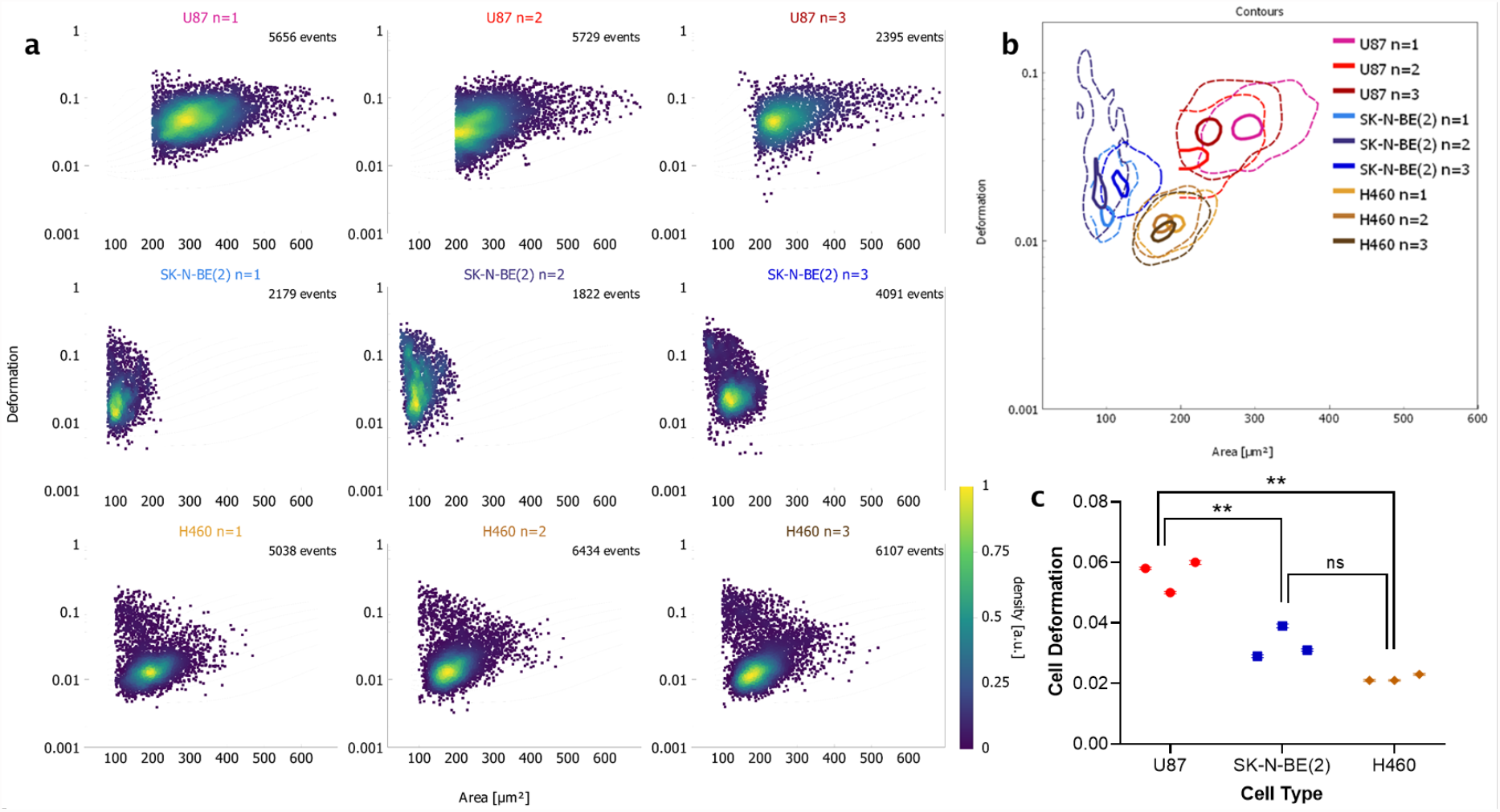
**Single cell deformation of U87, SK-N-BE(2) and H460 cells,** measured using force imaging cytometry (AcCellerator, Zellmechanik Dresden). Data from Supplementary Figure 10 was similarly used to calculate the degree of deformation caused by sheath flow (0.12 uL s^−1^). **(d)** Scatter plots generated of cell area versus cell deformation for each biological run. Heat map representative of count rate across single cell scatter. **(e)** Contour plots of deformation for each cell line. *Solid line,* 95^th^ percentile. *Dotted lines*, 50^th^ percentile of total cell population. The average of these values was then used in **(f)** to calculate significant differences of cell deformation potential between cell types. *Points*, mean of biological replicates. *Bars*, SEM. Significance by unpaired t-tests with Tukey correction, ns, non-significant, ** *p* < 0.01.

