## Supplementary figures for "Spatial-temporal analysis of nanoparticles in live tumor spheroids impacted by cell origin and density"

**SUPPLEMENTARY DATA**

### Deceased

**SUPPLEMENTARY DATA**

**Supplementary Tables**

**Supplementary Table 1: Cell characteristics of U87, SK-N-BE(2) and H460** with the mean and median diameters, alongside cell counts and deformation. Quantified using single-cell force imaging cytometry at a total flow rate of 0.160 uL $s^{-1}$ ........ 12

#### SUPPLEMENTARY DATA

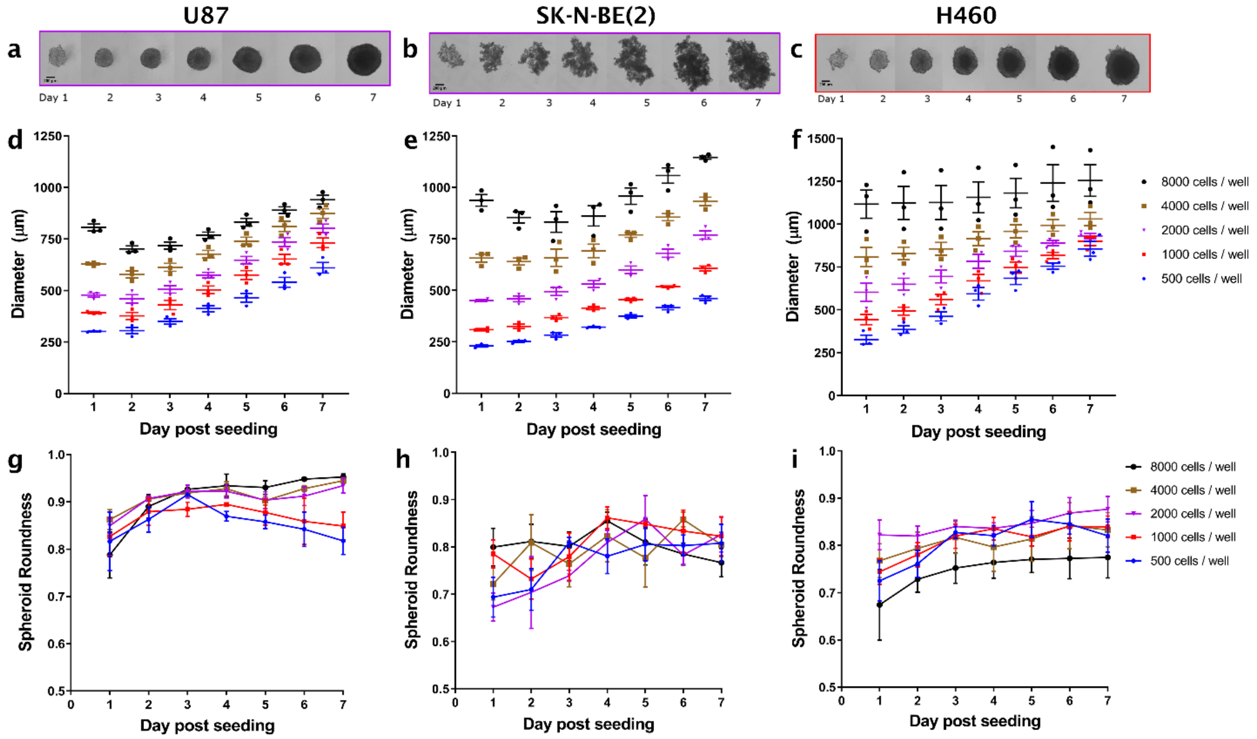

**Supplementary Figure 1: Spheroid growth of glioblastoma (U87), neuroblastoma (SK-N-BE(2)) and non-small cell lung cancer (H460)** seeded at Day 0 and imaged daily for 7 days. **(a)** Representative growth of U87, **(b)** SK-N-BE(2) spheroid (2000 cells at Day 0). **(c)** Representative growth of H460 spheroid (1000 cells at Day 0). Scale bar, 200 µm. This growth was quantified in ImageJ by measuring the diameter each day in **(d)** U87, **(e)** SK-N-BE(2) and **(f)** H460 spheroids at different seeding densities indicated in (f). Points, individual biological replicates (n=3) per time point, per seeding density. Lines, mean of n=3 ± SEM. Growth characteristics were quantified by aspect ratio, defined above as roundness in **(g)** U87, **(h)** SK-N-BE(2) and **(i)** H460 spheroids at the same seeding densities as above. For ease of visualisation, points here represent mean of n=3. Bars, SEM.

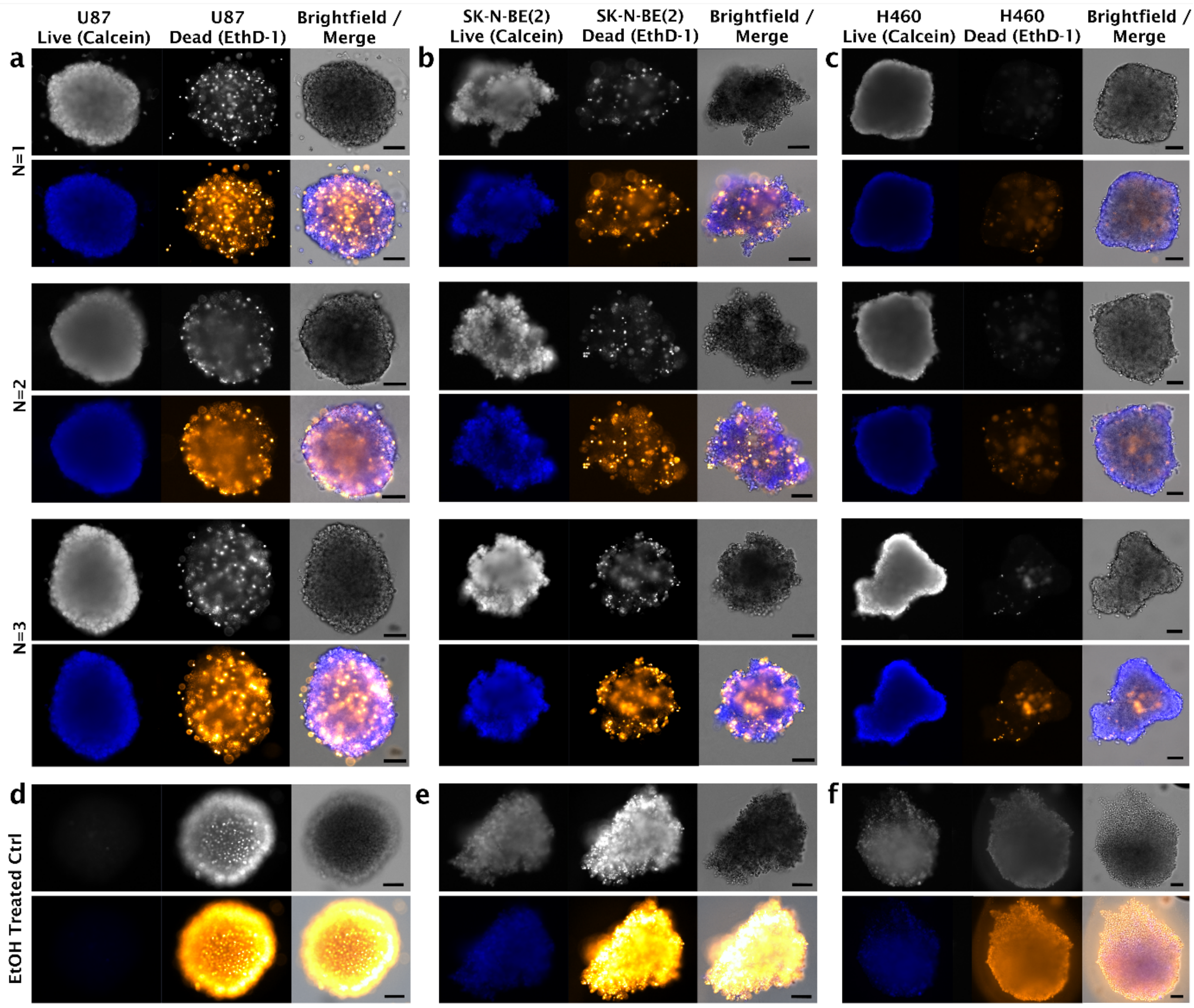

**Supplementary Figure 2: Spheroid viability and characteristics visualized through a live/dead assay** using Calcein and ethidium homodimer-1 (EthD-1). Representative brightfield and live (blue)/dead (orange-yellow) images of **(a)** U87 and **(b)** SK-N-BE(2) and **(c)** H460 cell spheroids, which were grown for 3 days in low adherent round-bottom well plates with an initial seeding density of $2 \times 10^3$, or $8 \times 10^2$ cells for H460 specifically. Viability is contrasted against ethanol (EtOH) treated controls for **(d)** U87 and **(e)** SK-N-BE(2) and **(f)** H460 respectively. Scale bar, 100 μm.

SUPPLEMENTARY DATA

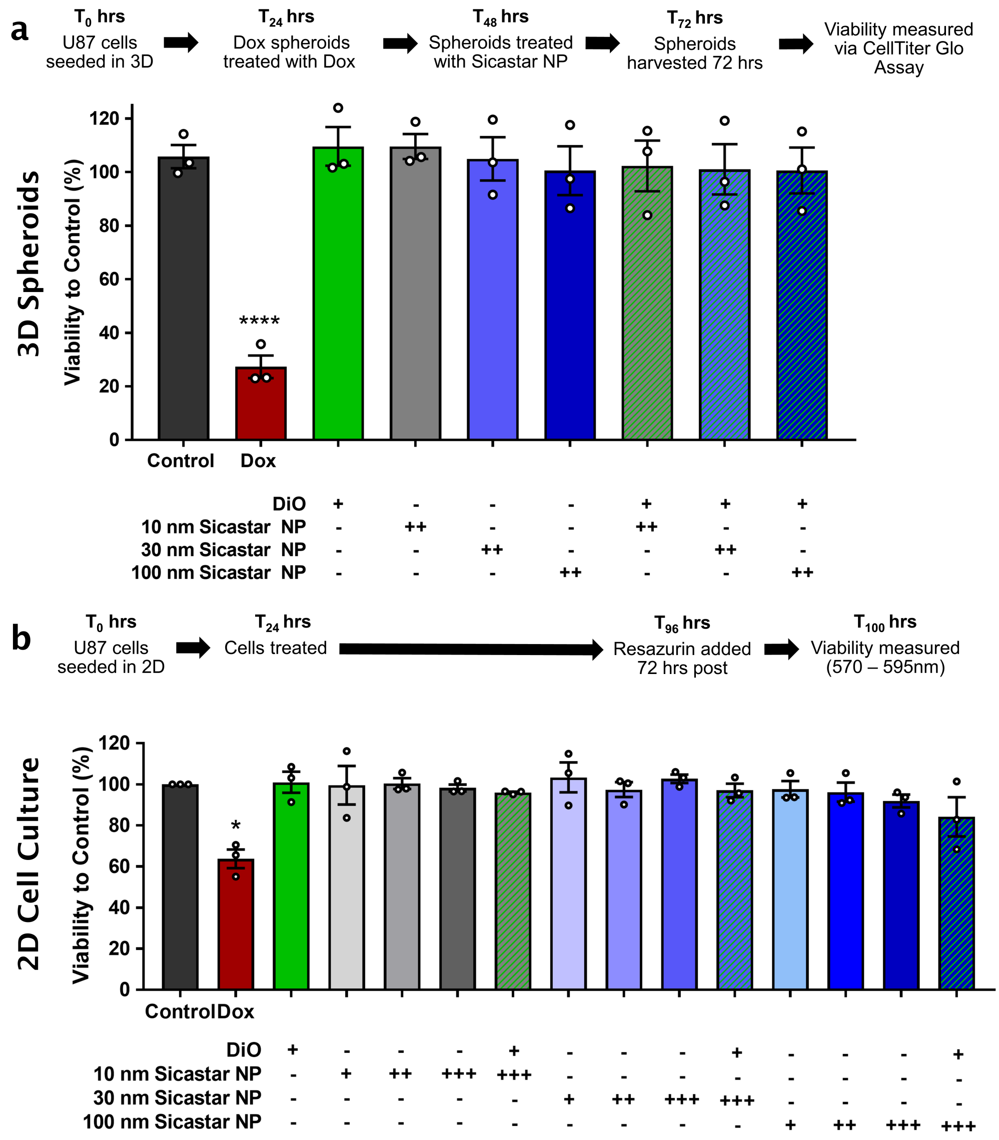

**Supplementary Figure 3: Cell viability of glioblastoma (U87) cells treated with silica nanoparticles (SiNPs) and membrane dye (DiO). (a)** 3D cell viability of U87 following treatment SiNPs alone or in combination with membrane dye (1 µM) for 24 hrs. Measured using Celltiter Glo Assay. Doxorubicin (Dox, 50 uM) used as a positive control. *Points*, biological replicates. *Columns,* mean of n = 3. *Bars,* SEM. Significance to control (untreated) using one-way ANOVA, **** $p < 0.0001$. **(b)** 2D cell viability of U87 following treatment with SiNP (+ 1 ug / mL, ++ 10 ug / mL, +++ 100 ug / mL, as indicated) membrane dye (1 µM) and combination over 72 hours. Doxorubicin (Dox, 0.25 uM) used as a positive control. Measured using a Resazurin-based cell viability assay. *Points*, biological replicates. *Columns,* mean of n = 3. *Bars,* SEM. Significance to control (untreated) using a paired ratio t-test, * $p < 0.05$.

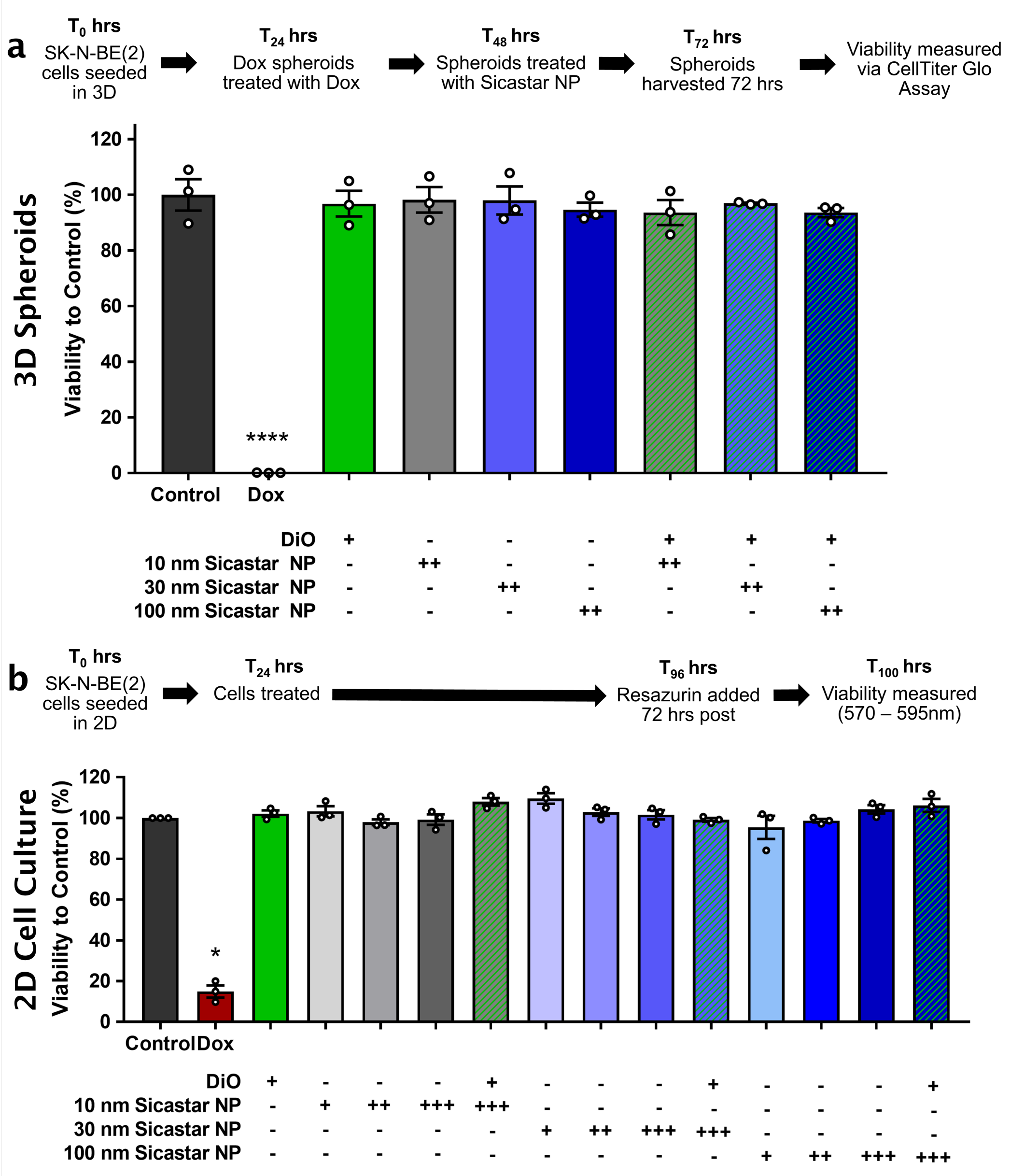

**Supplementary Figure 4: Cell viability of neuroblastoma (SK-N-BE(2)) cells treated with silica nanoparticles (SiNPs) and membrane dye (DiO). (a)** 3D cell viability of SK-N-BE(2) following treatment SiNPs alone or in combination with membrane dye (1 µM) for 24 hrs. Measured using Celltiter Glo Assay. Doxorubicin (Dox, 50 uM) used as a positive control. *Points*, biological replicates. *Columns,* mean of n = 3. *Bars,* SEM. Significance to control (untreated) using one-way ANOVA, **** $p < 0.0001$. **(b)** 2D cell viability of SK-N-BE(2) following treatment with SiNP (+ 1 ug / mL, ++ 10 ug / mL, +++ 100 ug / mL, as indicated) membrane dye (1 µM) and combination over 72 hours. Doxorubicin (Dox, 0.25 uM) used as a positive control Measured using a Resazurin-based cell viability assay. *Points*, biological replicates. *Columns,* mean of n = 3. *Bars,* SEM. Significance to control (untreated) using a paired ratio t-test, * $p < 0.05$.

### SUPPLEMENTARY DATA

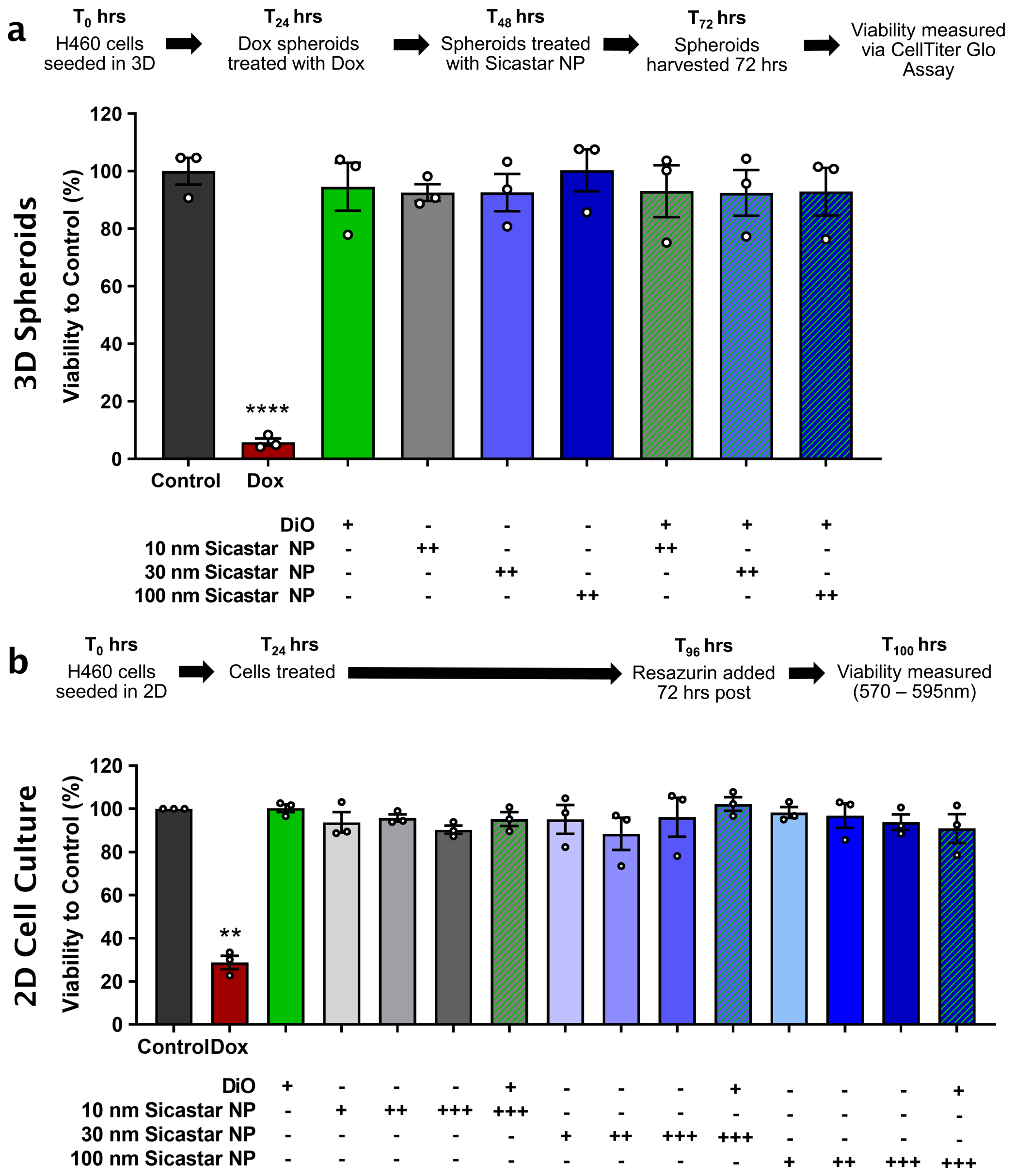

**Supplementary Figure 5: Cell viability of non-small cell lung cancer (H460) cells treated with silica nanoparticles (SiNPs) and membrane dye (DiO). (a)** 3D cell viability of H460 following treatment SiNPs alone or in combination with membrane dye (1 µM) for 24 hrs. Measured using Celltiter Glo Assay. Doxorubicin (Dox, 20 uM) used as a positive control. *Points*, biological replicates. *Columns,* mean of n = 3. *Bars,* SEM. Significance to control (untreated) using one-way ANOVA, **** $p < 0.0001$. **(b)** 2D cell viability of H460 following treatment with SiNP (+ 1 ug / mL, ++ 10 ug / mL, +++ 100 ug / mL, as indicated) membrane dye (1 µM) and combination over 72 hours. Doxorubicin (Dox, 0.1 uM) used as a positive control. Measured using a Resazurin-based cell viability assay. *Points*, biological replicates. *Columns,* mean of n = 3. *Bars,* SEM. Significance to control (untreated) using a paired ratio t-test, ** $p < 0.01$.

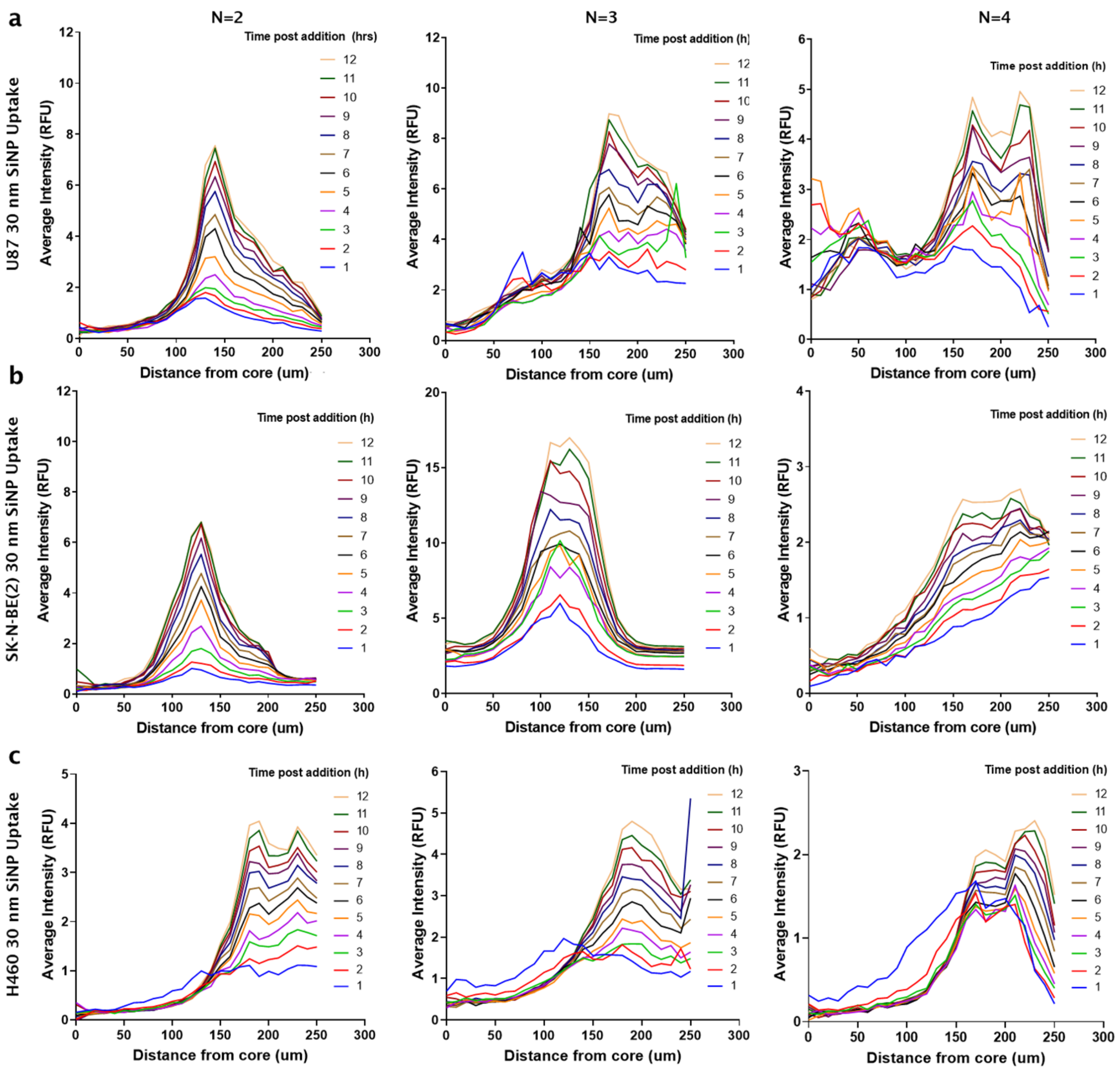

**Supplementary Figure 6: 3D Azimuthal quantification of 30 nm SiNP uptake** in **(a)** glioblastoma (U87), **(b)** neuroblastoma (SK-N-BE(2)) and **(c)** non-small cell lung cancer (H460) cell spheroids. Each graph represents 30 nm SiNP tumor spheroid uptake of a biologically independent experiment, in addition to the data presented elsewhere.

#### SUPPLEMENTARY DATA

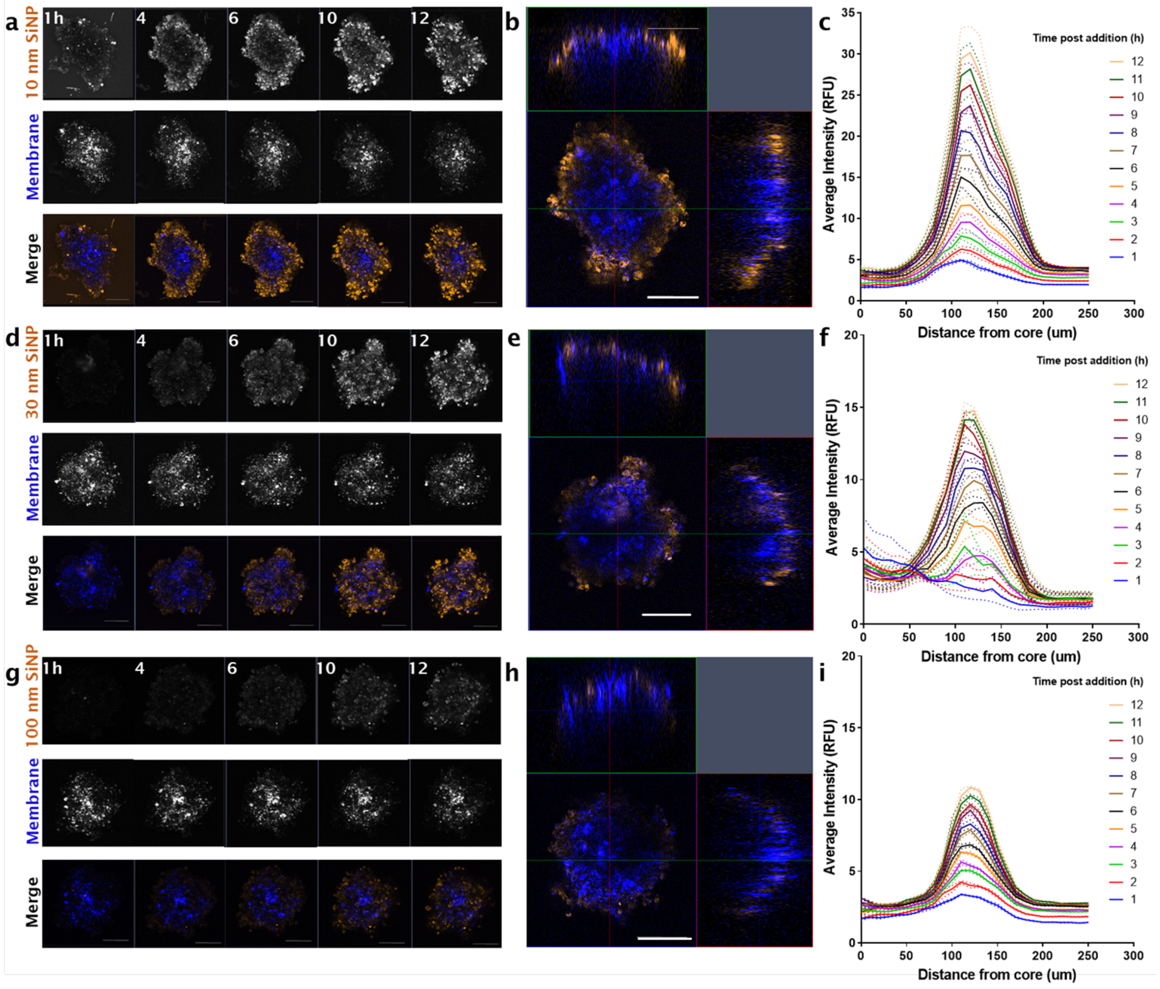

**Supplementary Figure 7: Silica nanoparticle (SiNP) uptake in neuroblastoma (SK-N-BE(2)) tumor spheroids over 12 hours**. **(a)** Representative maximum intensity projections of 10 nm SiNP, membrane (DiO) and merge at 1, 4, 6, 10, 12 hours post addition (Orange, SiNP; Blue, Membrane). Z-stack images acquired using a Zeiss 880 confocal microscope (Fast Airy, sequential frame-fast laser excitation at 488 nm and 561 nm, 10X objective). **(b)** Orthogonal (XY, XZ, ZY) merge of SK-N-BE(2) spheroid at six hours post SiNP addition, representative of n = 4. **(c)** Representative quantification of nanoparticle uptake from the core of the spheroid to the circumference over time with increased 10 nm SiNP penetration. Analysis conducted using a 3D azimuth averaging custom script, MATLAB (2020a). Workflow above was performed for 30 nm SiNP showing **(d)** maximum intensity projections over 1, 4, 6, 10, 12 hours post SiNP addition; **(e)** orthogonal merge at six hours and **(f)** azimuth quantification respectively. Imaging and analysis also performed for 100 nm SiNP in panels **(g)** maximum intensity projections; **(h)** orthogonal merge at six hours and **(i)** quantification of 100 nm SiNP uptake. Lines, mean of t=3 analysis iterations. Dotted lines, error SEM. Scale bar, 100 μm.

SUPPLEMENTARY DATA

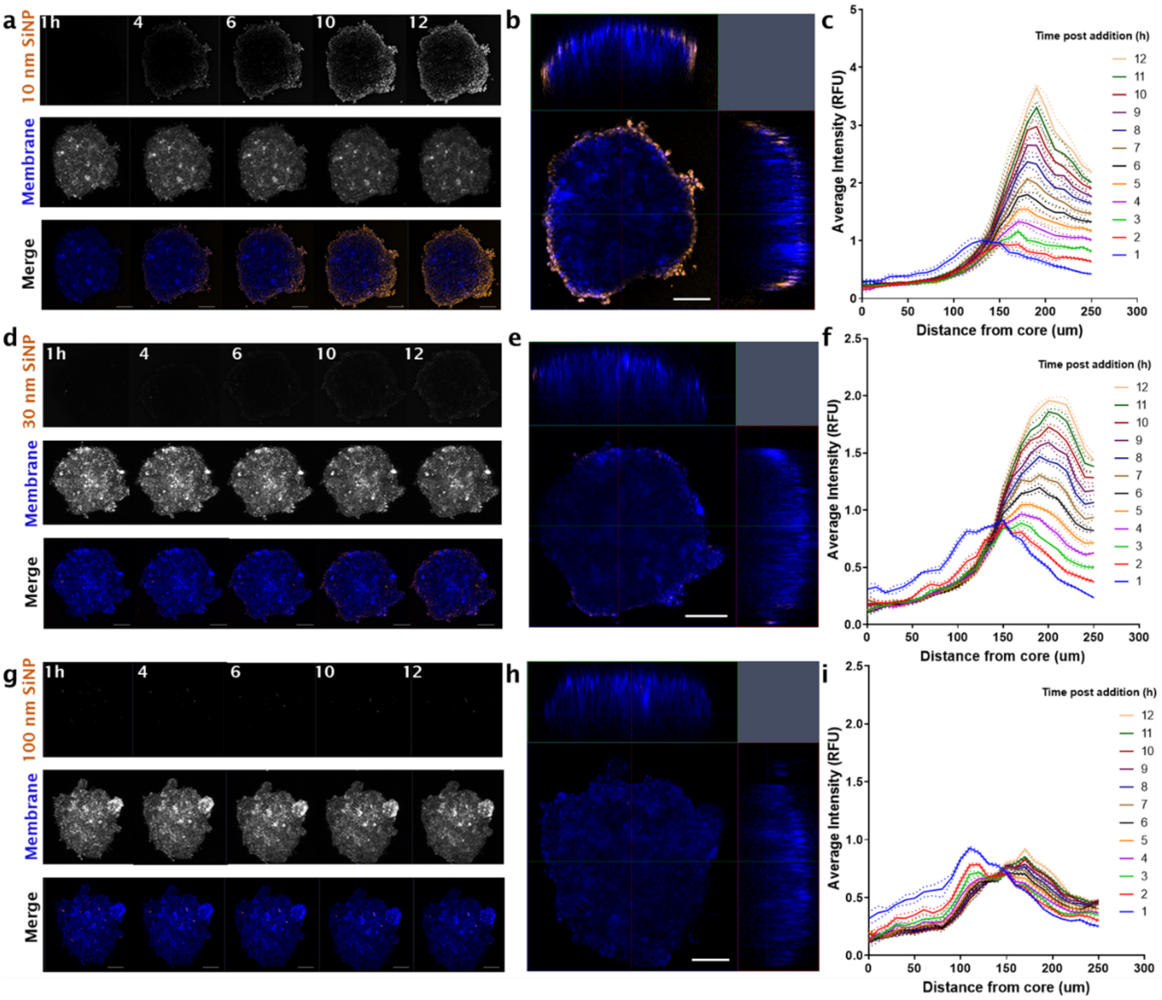

**Supplementary Figure 8: Silica nanoparticle (SiNP) uptake in non-small cell lung cancer (H460) tumor spheroids over 12 hours**. **(a)** Representative maximum intensity projections of 10 nm SiNP, membrane (DiO) and merge at 1, 4, 6, 10, 12 hours post addition (Orange, SiNP; Blue, Membrane). Z-stack images acquired using a Zeiss 880 confocal microscope (Fast Airy, sequential frame-fast laser excitation at 488 nm and 561 nm, 10X objective). **(b)** Orthogonal (XY, XZ, ZY) merge of H460 spheroid at six hours post SiNP addition, representative of n = 4. **(c)** Representative quantification of nanoparticle uptake from the core of the spheroid to the circumference over time with increased 10 nm SiNP penetration. Analysis conducted using a 3D azimuth averaging custom script, MATLAB (2020a). Workflow above was performed for 30 nm SiNP showing **(d)** maximum intensity projections over 1, 4, 6, 10, 12 hours post SiNP addition; **(e)** orthogonal merge at six hours and **(f)** azimuth quantification respectively. Imaging and analysis also performed for 100 nm SiNP in panels **(g)** maximum intensity projections; **(h)** orthogonal merge at six hours and **(i)** quantification of 100 nm SiNP uptake. Lines, mean of t=3 analysis iterations. Dotted lines, error SEM. Scale bar, 100 µm.

SUPPLEMENTARY DATA

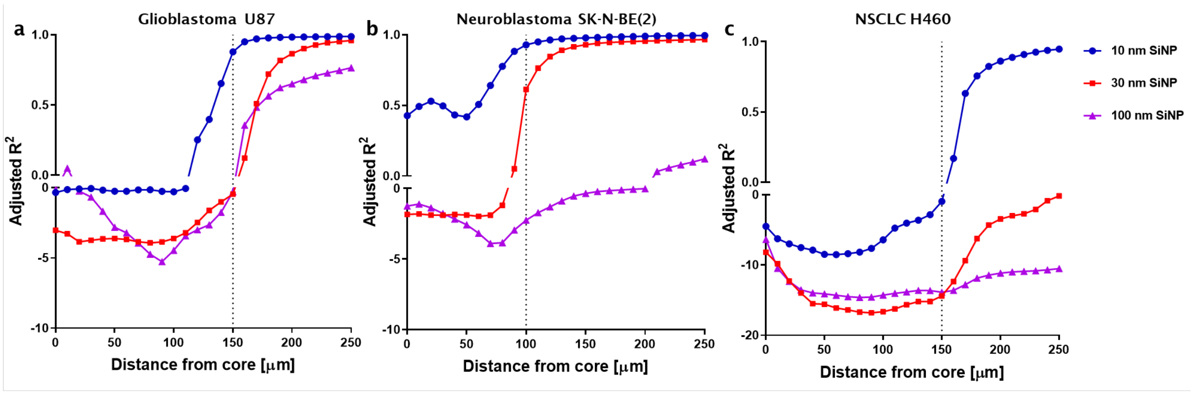

**Supplementary Figure 9: Adjusted r squared ($R^2$) fit values for silica nanoparticle (SiNP) diffusion kinetics** for **(a)** glioblastoma U87 **(b)** neuroblastoma (SK-N-BE(2)) and **(c)** H460 spheroids, calculated using the forward in time, central in space (FTCS) coefficient method. Dotted lines, right side indicating improved fit of model with $R^2 \rightarrow 0.99$, excluding 100 nm SiNP in SK-N-BE(2) and 30 nm, 100 nm in H460 which did not show $R^2$ above 0.25.

**Supplementary Table 1: Cell characteristics of U87, SK-N-BE(2) and H460** with the mean and median diameters, alongside cell counts and deformation. Quantified using single-cell force imaging cytometry at a total flow rate of 0.160 uL $s^{-1}$.

| Cell Line | Diameter (µm) Mean ± SD | Diameter (µm) Median ± SD | Deformation Mean ± SD | Cell Count ($n_{total}$) |
|---|---|---|---|---|
| U87 | 19.61 ± 0.54 | 19.05 ± 0.66 | 0.056 ± 0.004 | 13663 |
| SK-N-BE(2) | 12.49 ± 0.45 | 12.16 ± 0.54 | 0.033 ± 0.004 | 7160 |
| H460 | 15.93 ± 0.08 | 15.64 ± 0.11 | 0.022 ± 0.001 | 17336 |

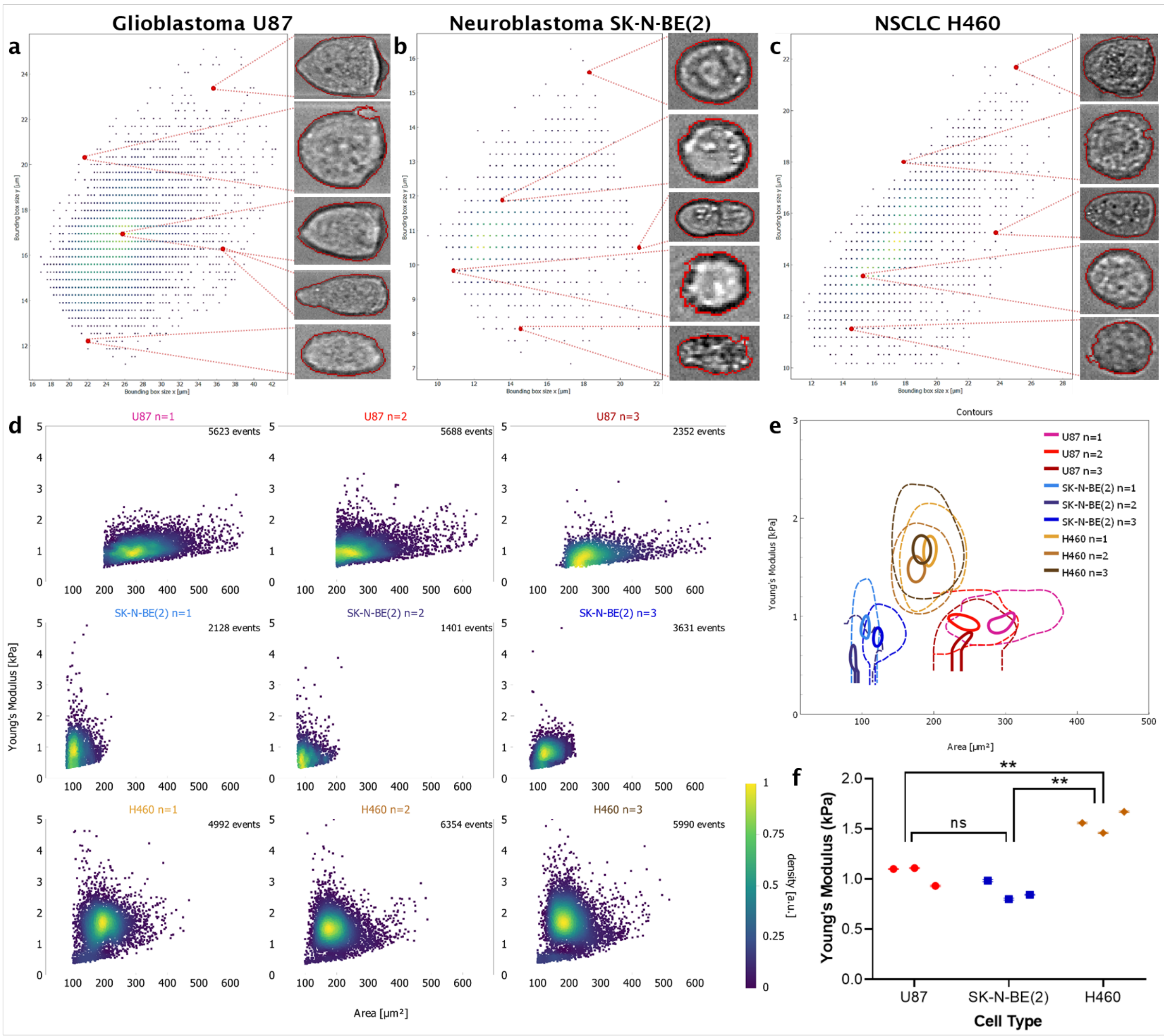

**Supplementary Figure 10: Single cell stiffness of U87, SK-N-BE(2) and H460 cells,** measured using force imaging cytometry (AcCellerator, Zellmechanik Dresden). Representative images of single cells of **(a)** U87, **(b)** SK-N-BE(2) and **(c)** H460 which were imaged at a total flow rate of 0.16 uL $s^{-1}$. Cells were gated according to size in X and Y as above to ensure single cell suspensions. **(d)** Scatter plots generated of cell area versus Young's Modulus calculated for each biological run, 1400 minimum cells per run. Heat map representative of count rate across single cell scatter. **(e)** Contour plots of Young's Modulus for each cell line. *Solid line,* 95$^{th}$ percentile. *Dotted lines*, 50$^{th}$ percentile of total cell population. The average of these values was then used in **(f)** to calculate significant differences of Young's Modulus between cell types. *Points*, mean of biological replicate. *Bars*, SEM. Significance by unpaired t-tests with Tukey correction, ns, non-significant, ** $p < 0.01$.

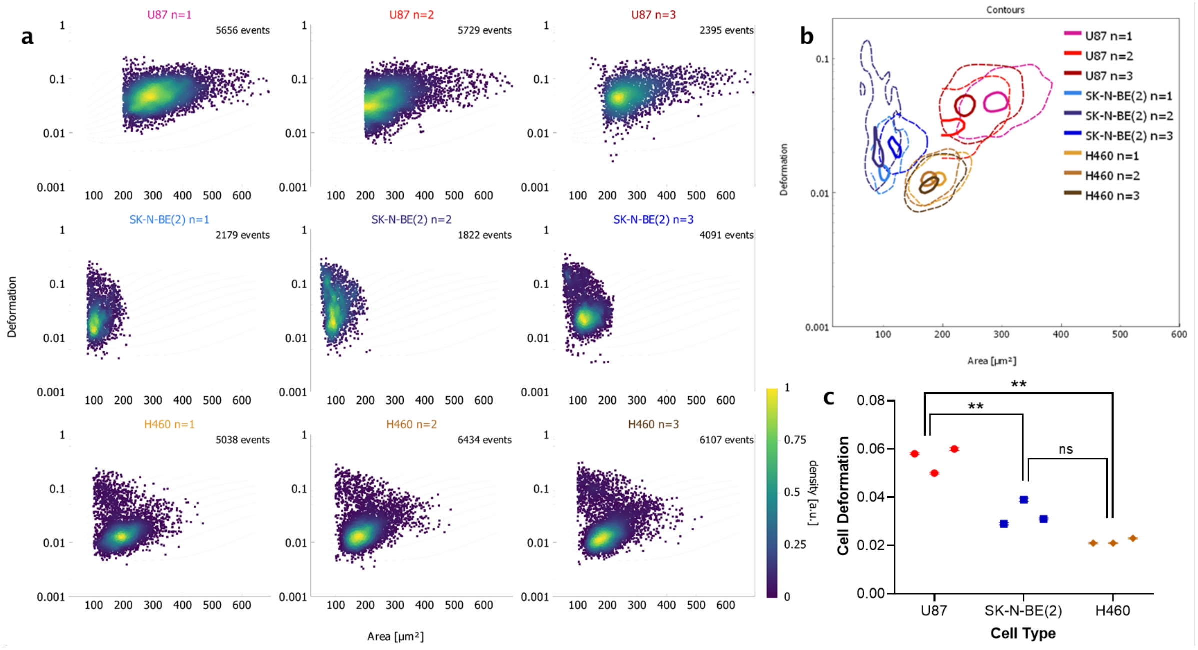

**Supplementary Figure 11: Single cell deformation of U87, SK-N-BE(2) and H460 cells,** measured using force imaging cytometry (AcCellerator, Zellmechanik Dresden). Data from Supplementary Figure 10 was similarly used to calculate the degree of deformation caused by sheath flow (0.12 uL $s^{-1}$). **(d)** Scatter plots generated of cell area versus cell deformation for each biological run. Heat map representative of count rate across single cell scatter. **(e)** Contour plots of deformation for each cell line. *Solid line,* 95$^{th}$ percentile. *Dotted lines*, 50$^{th}$ percentile of total cell population. The average of these values was then used in **(f)** to calculate significant differences of cell deformation potential between cell types. *Points*, mean of biological replicates. *Bars*, SEM. Significance by unpaired t-tests with Tukey correction, ns, non-significant, ** $p < 0.01$.
